# Environmental Demands Outweigh Age Effects in Cortical Dynamics During Walking

**DOI:** 10.64898/2026.09.19.751869

**Authors:** Danishta Kaul, Magda Mustile, Mark Donoghue, Alexander Brownlee, Magdalena Ietswaart, Gemma Learmonth

## Abstract

Reduced visibility is a common contributor to falls in older adults, but the neural processes that support safe locomotion under low lighting are not well established. In this study, 22 young (mean age = 22.5) and 20 older (mean age = 70.9) adults walked along a pathway under bright, ambient and dark lighting conditions while stepping over expected and unexpected obstacles projected on the floor. Mobile electroencephalography (EEG), combined with motion capture, was used to quantify spectral power changes during obstacle crossing preparation and the post-crossing phases, relative to a standing baseline. We identified that reduced lighting conditions elicited a widespread increase in cortical engagement, predominantly in the alpha band, during both preparation and reset phases. Obstacle negotiation recruited a distributed sensorimotor network, with reduced theta-alpha-beta desynchronisation during preparation and a robust post-movement beta rebound during the post-crossing reset phase. Expectation effects were modest, with unexpected obstacles inducing greater transient alpha-beta desynchronisation relative to expected obstacles, suggesting additional cortical resources are recruited under uncertainty. Age effects were minimal and limited to reduced frontal alpha desynchronisation in older adults compared to young adults during the preparation phase, possibly reflecting differences in walking speed rather than differences in neural control. In conclusion, environmental features such as low lighting and obstacle presence exert a stronger influence on cortical dynamics during walking than age and expectation effects alone. This provides a foundation for future work examining how these processes may differ in older adults at increased risk of falls.

## 1. Introduction

As the global population ages, addressing the challenges faced by older adults in activities of daily living has become an increasing public health priority (Kwon et al., 2023). Among the challenges associated with ageing, falls resulting from slipping and tripping during walking are a leading public health concern (Luo et al., 2021), with one-third of adults aged over 65 affected by falls annually (Ambrose et al., 2013). In Scotland, falls are the primary reason for ambulance callouts to assist older people, and over 18,000 older people are hospitalised each year after falling (NHS inform, 2025). These incidents have serious consequences for both the physical and mental well-being of older people, including injury, decreased physical activity, functional decline, fear of falling, anxiety, and depressive symptoms (Jacob et al., 2023; Vieira et al., 2016). Moreover, falls impose a substantial economic burden on healthcare systems, costing the NHS approximately £630 million per year (Boot et al., 2023).

There is a critical need to characterise the sensory and environmental factors that contribute to fall risk in older adults. Postural control and locomotion depend on the coordinated integration of visual, vestibular, and somatosensory inputs (Horak, 2006; Kolbaşı et al., 2025). Within this multisensory system, visual functions such as acuity, contrast sensitivity, and depth perception support obstacle detection and distance estimation, and age-related declines in these visual processes are strongly associated with increased fall risk in older adults (Alghwiri & Whitney, 2020). Scotopic vision is particularly vulnerable to ageing, and older adults frequently report difficulty with night-time vision and activities such as driving at night (Owsley, 2016; Rundus et al., 2025). Psychophysical studies demonstrate that ageing is associated with reduced light sensitivity in dark conditions, even after full dark adaptation (Jackson & Owsley, 2000; Owsley, 2016; Sturr et al., 1997), and that this decline in sensitivity is greater than age-related losses in photopic (daytime) vision (Jackson & Owsley, 2000; Owsley, 2016).

Beyond optical factors such as lens density and pupillary miosis, which partly account for elevated scotopic thresholds in older adults (i.e., the lowest level of light that can be detected after full dark adaptation; Grewal et al., 2021), scotopic dysfunction has also been linked to age-related changes in the visual cycle, particularly slower rhodopsin regeneration, which prolongs dark adaptation (Bakdalieh et al., 2025; Owsley, 2016). These delays have clear functional consequences. For example, it takes individuals in their seventies 10 minutes longer than young adults to recover pre-bleach sensitivity following bright light exposure (bleaching occurs when bright light converts visual pigment to a colourless form, temporarily reducing visual sensitivity; Lamb & Pugh, 2004), increasing the likelihood of errors during transitions from bright to dim lighting conditions (Owsley, 2016). Such delays are particularly relevant in everyday environments where lighting changes abruptly and may contribute to a heightened risk of falls.

Lighting is a persistent feature of the physical environment and plays a critical role in mobility and safety. Inadequate lighting has been identified as a key environmental barrier, limiting older adults’ engagement in their physical environment and increasing risk of falls (Lu et al., 2019). Reduced lighting alters gait patterns in older adults, often resulting in slower walking speeds and more cautious movement strategies. For example, Moe-Nilssen et al. (2006) showed that abrupt transitions from normal to dim lighting challenged balance during walking in older women, leading to protective gait patterns such as shorter steps and higher cadence. Other studies have demonstrated that providing visual cues in low-light environments can improve gait performance in older adults. Figueiro et al. (2011) showed that augmenting dim night lighting with laser lines outlining the pathway improved walking speed and reduced step length variability in older adults at high risk of falling. Similarly, Luo et al. (2022) found that LED lighting used to increase the salience of the walking destination improved older adults’ gait speed and walking stability. Age-related differences in gait under low illumination have also been observed. Naaman et al. (2023) found that during treadmill walking young adults’ gait stability was largely unaffected by darkness, whereas older adults increased their stride time variability in the dark, suggesting a greater reliance on vision to maintain gait stability in older adults.

Despite the role of vision and lighting in gait and balance, the neural processes that support locomotion under reduced visual input are poorly understood. Traditional neuroimaging methods such as fMRI and MEG constrained participants’ movements to minimise motion-related artifacts, preventing the investigation of walking under naturalistic conditions and limiting the understanding of brain activity during real-world locomotion (Richer et al., 2024). However, recent technological advancements in mobile electroencephalography (EEG) have broadened the range of environments in which brain signals can be studied (Ladouce et al., 2019; Mustile et al., 2021; Park et al., 2018).

Cao et al. (2020) investigated cortical alpha activity during overground walking under light and dark conditions, with the dark condition created using a blindfold and opaque covering. Occipital alpha power decreased during walking relative to standing in both conditions, indicating that walking-related alpha changes occur regardless of visual input. As alpha power in sensory areas is associated with inhibition of sensory processing (Cao et al., 2020; Klimesch et al., 2007), the walking-related decrease was interpreted as a shift in attentional state in which peripheral visual input became less suppressed. This interpretation is supported by evidence that walking increases the processing of peripheral visual information, which plays a key role in guiding movement and navigation (Cao & Händel, 2019). Occipital alpha power was also lower in the light than in the dark, further supporting the interpretation that alpha activity reflects visual processing.

Oliveira et al. (2017) examined EEG activity during treadmill walking with eyes open and eyes closed. The study reported that, compared to walking with eyes open, the eyes closed condition produced stronger theta desynchronisation over frontal and premotor cortices during stance, and stronger desynchronisation from theta to beta band in bilateral somatosensory cortex during the shift to single-support phase. The authors interpreted the stronger theta to beta desynchronisation as reflecting greater engagement of the somatosensory cortices in movement control, particularly for sustaining balance while walking. This conclusion is supported by findings from Sipp et al. (2013), who reported decreased beta power in sensorimotor areas when walking on a balance beam, consistent with the notion that motor control is associated with a reduction in beta power (Engel & Fries, 2010; Pfurtscheller et al., 1996). Overall, the findings indicate that when vision is unavailable, the brain engages in sensory reweighting, reflected in gait-phase specific changes in cortical activity in regions involved in sensory processing and integration (Oliveira et al., 2017).

Beyond the effects of restricted vision, walking itself may require greater cortical engagement in older adults than younger adults (Salminen et al., 2025). Salminen et al. (2025) found that theta power in the left posterior parietal region increased more with faster walking speed in older than in younger adults, suggesting that older adults may rely more on cortical resources for gait control at faster speeds due to reduced automaticity of walking with ageing.

Given that tripping, particularly during obstacle negotiation, is one of the most frequently cited causes of falls (Cho et al., 2013; Wang et al., 2020), it is important to understand the cognitive processes supporting ambulatory motor control. The Dual Mechanisms of Control framework distinguishes between proactive control, which involves anticipating and preventing interference before it has happened, and reactive control, which focuses on detecting and resolving interference after it has started (Braver, 2012). During obstacle negotiation, proactive control allows gait to be adjusted in advance of an upcoming obstacle, while reactive control supports recovery when balance is unexpectedly perturbed (Mustile et al., 2021). This dual-mode framework can be used to understand the electrocortical signatures observed in studies on obstacle navigation. For instance, Nordin et al. (2019) recorded EEG activity while young adults navigated unexpected obstacles on a treadmill. Within approximately 200 ms of obstacle appearance, power increased across the delta, theta and alpha bands (3–13 Hz) over the supplementary motor area and premotor cortex, followed by a later increase over the posterior parietal cortex prior to obstacle crossing. The authors proposed that the supplementary motor area and premotor cortex were involved in disrupting the gait cycle in response to the obstacle, while the posterior parietal cortex supported preparing foot placement for obstacle crossing.

A study by Mustile et al. (2021) identified the neural markers of proactive and reactive control using mobile EEG. Frontal theta power increased more before crossing unexpected obstacles compared to expected obstacles or obstacle-free walking, with the increase coinciding with the appearance of the unexpected obstacle. This was interpreted as a marker of proactive control, engaged when participants had less time to prepare to navigate the obstacle. Compared to obstacle-free walking, beta power decreased more over sensorimotor areas as participants approached the obstacle, reflecting heightened motor readiness. After obstacle crossing, a greater increase in parietal beta power was observed, indicating a beta rebound, which the authors interpreted as a reactive control signature reflecting resetting of the motor system.

Sensory information is also important for navigating the environment. Vision underpins the proactive control of locomotion, allowing potential disruptions to balance to be identified and avoided in advance (Patla, 1997). Consequently, low-light conditions reduce the visual information needed for obstacle detection and the regulation of body stability, leading to a heightened risk of falling in older adults (Cho et al., 2013; Lu et al., 2019). Moreover, tripping over obstacles is a key contributor to falls (Cho et al., 2013). However, although studies have demonstrated that restricted vision alters cortical activity during walking (Cao et al., 2020; Oliveira et al., 2017), these studies did not involve obstacle navigation. Conversely, studies that have examined the neural correlates of obstacle navigation have done so only under well-lit conditions (Mustile et al., 2021; Nordin et al., 2019). To date, no study has examined how different lighting conditions influence neural activity during obstacle navigation. Furthermore, although age-related declines in scotopic sensitivity and delays in dark adaptation are well-documented (Owsley, 2016), and older adults are at increased risk of falling when negotiating obstacles under low lighting (Cho et al., 2013), the neural mechanisms underlying these age-related difficulties remain unexplored.

Thus, the present study investigates how different light conditions influence walking behaviour and obstacle negotiation in young and older adults. Participants walked along a track where obstacles were projected either before they started walking (Expected Present) or after they had started walking (Unexpected Present), under 3 different lighting conditions: a well-lit condition with room lighting turned on, an ambient condition with dim light provided by LEDs, and a dark condition with the lights turned off. Mobile EEG was recorded to identify the neural dynamics of obstacle negotiation during naturalistic walking. Given age-related declines in contrast sensitivity and delayed dark adaptation, contrast sensitivity was measured under each light condition, and participants were given sufficient time to adapt before they started walking. This approach aimed to capture realistic visual challenges encountered in everyday environments and to clarify how ageing and lighting interact to influence behavioural and neural mechanisms of locomotion.

## 2. Methods

### 2.1. Participants

A total of 44 participants were recruited (22 young adults aged 18–30, and 22 older adults aged 65–80). For the EEG analysis, data from two older adults were excluded: one due to excessive EEG artifacts, and one due to insufficient trials remaining after preprocessing. The final EEG sample therefore consisted of 42 participants (N_young_ = 22, mean age = 22.5 years, SD = 3.3; N_older_ = 20, mean age = 70.9 years, SD = 3.5). The young group included 15 women, 6 men, and 1 non-binary participant, with 18 self-reporting as right-handed, 3 ambidextrous, and 1 left-handed. The older group included 10 women and 10 men, with 16 right-handed, 2 ambidextrous, and 2 left-handed. For the gait analysis, one additional older adult was excluded due to insufficient trials remaining following the removal of trials with missing marker data. The final sample for gait analysis consisted of 41 participants (N_young_ = 22, mean age = 22.5 years, SD = 3.3; N_older_ = 19, mean age = 70.7 years, SD = 3.5).

Participants were eligible to take part if they had no known neurological disorders or problems with walking, and normal or corrected-to-normal vision. The sample size was determined by considering both expected feasibility, and prior studies using mobile EEG with 32 to 40 participants (Mustile et al., 2021; Vila-Chã et al., 2022). Ethical approval was granted by the University of Stirling General University Ethics Panel. All participants provided written informed consent prior to participation.

Self-reported physical activity was assessed using a checklist based on the National Health Service (National Health Service, n.d.) and UK Chief Medical Officers’ physical activity guidelines (Department of Health and Social Care, 2026). 91% of young participants and 95% of older participants reported meeting at least one recommended activity threshold. Two young and one older participant reported engaging only in light physical activity.

### 2.2. Procedure and materials

The study used a mixed design with 3 variables: Age (Young and Older), Lighting (Bright, Ambient, and Dark) and Obstacle (Expected Present, Unexpected Present, Expected Absent, and Unexpected Absent).

First, participants were fitted with a mobile EEG cap connected to a lightweight backpack-mounted amplifier and lower-body kinematics were recorded using an OptiTrack motion capture system (Corvallis, OR, USA) with reflective markers placed according to the Rizzoli lower-body marker set (Leardini et al., 2007). Participants were instructed to walk back and forth along a 6.75 m carpeted walkway while stepping over 2D blue rectangular obstacles projected onto the floor. Approximately 67 cm at each end of the walkway was used as turnaround space, resulting in a walking distance of 5.41 m. Before the expected obstacle conditions, participants were instructed that obstacles, when present, would be visible before they started walking and that some trials would contain no obstacle. Before the unexpected obstacle conditions, participants were informed that an obstacle might or might not appear after they had started walking. Accordingly, in ‘Expected Present’ trials, the obstacle was visible before participants started walking, whereas in ‘Expected Absent’ trials no obstacle was visible before walking began. In ‘Unexpected Present’ trials, the obstacle appeared after participants had started walking, whereas in ‘Unexpected Absent’ trials no obstacle appeared (Figure 1).

**Figure 1.**
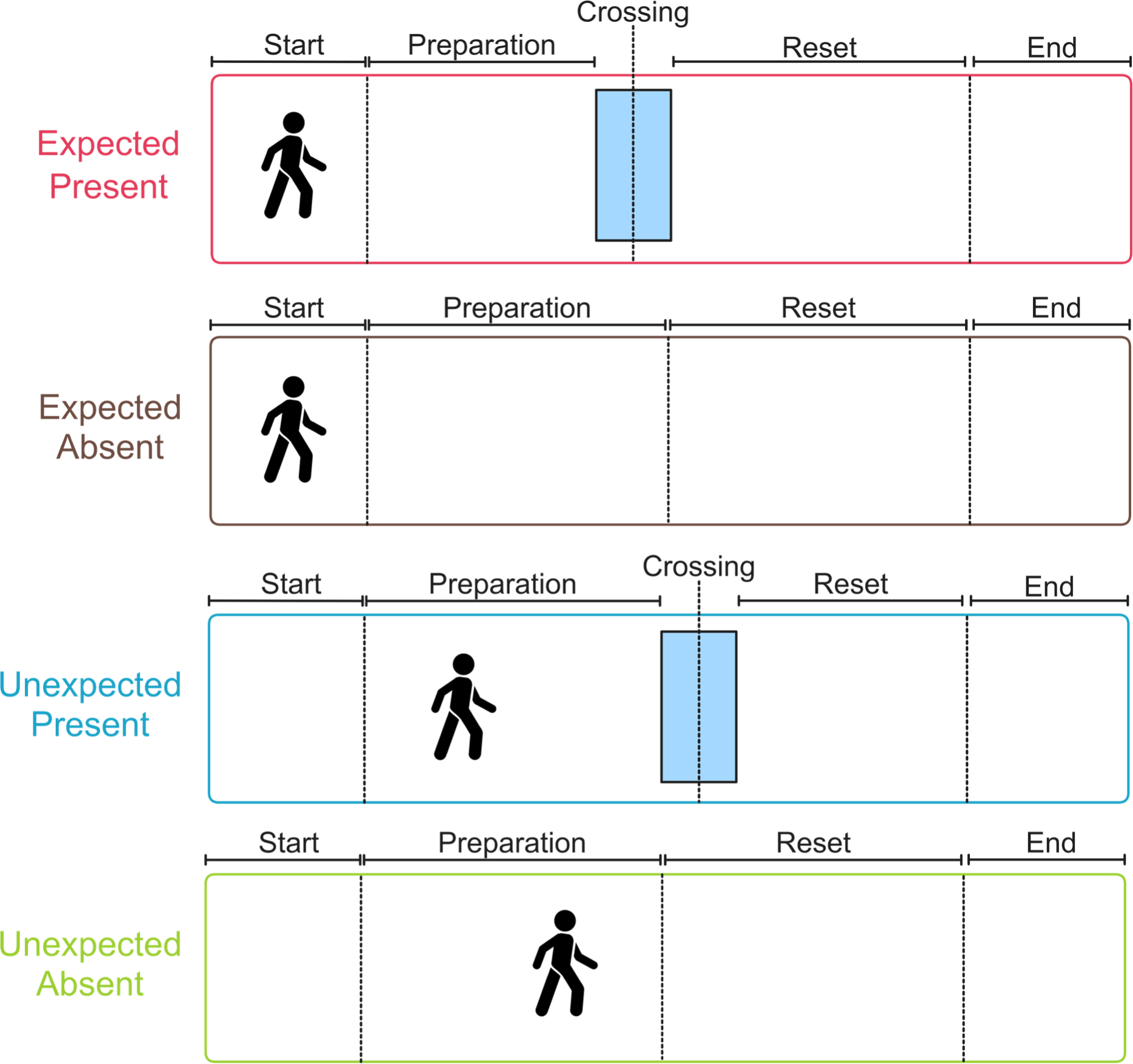
An illustration of the 4 obstacle conditions. In the Expected Present and Unexpected Present conditions, the preparation and reset phases are marked before and after the obstacle crossing event, respectively. In the Expected Absent and Unexpected Absent conditions, the preparation and reset phases are marked before and after reaching the mid-point of the walking path, respectively.

Each crossing of the walkway constituted one trial. At the end of each trial, a beep signalled participants to stop walking, turn to face the opposite end of the walkway, stand still while silently counting to 2 seconds, and then begin the next trial at a self-determined comfortable walking pace. This sequence was repeated for all trials. The 2 second standing period served as the baseline for EEG analysis.

For the ‘Expected Present’ and ‘Unexpected Present’ trials, obstacles appeared when the participant was either 1.2 m (near) or 2 m (far) away from the obstacle’s position. On each trial, to minimise expectation effects, the selected distance (near or far) was randomly jittered by ± 13 cm along the path. Additionally, in half of the trials, the obstacle width was proportional to the participant’s step length (see Supplementary Section S1), whereas in the remaining trials the width was increased by 18%.

Participants completed all trials under 3 different lighting conditions. In the Bright condition (12.82 cd/m²), the artificial lights in the room were turned on. The Ambient condition involved the use of dim LED lighting (0.162 cd/m²). In the Dark condition (<0.01 cd/m²), the lights were turned off. The order of the lighting blocks was counterbalanced across participants. Within each lighting block, the order of the expected and unexpected obstacle conditions was randomised. Within each obstacle condition, obstacle-present and obstacle-absent trials were presented in a random order.

The experiment was divided into 3 blocks, each with a different light condition (Figure 2). There were 80 trials in each block (240 trials in total), with 20 trials for each of the 4 obstacle conditions. Before beginning each block, participants were given 5 minutes to pre-adapt to the lighting condition of that block (which also served as a break), followed by an assessment of their contrast sensitivity. In step 1 of this assessment, participants were presented with 32 shades of the blue obstacle, ranging from the dimmest to the brightest blue, and were asked to identify the shade where the obstacle was barely visible. In step 2, the same 32 shades were presented again, but in reverse order - from the brightest to the dimmest blue - and participants were again asked to indicate the shade where the obstacle was barely visible. The mean response from step 1 and step 2 determined the initial barely perceptible shade. The obstacle was presented to the participants again in this shade, and the brightness was gradually lowered until they reported that they could no longer see the obstacle. The difference between the average shade and the shade where the obstacle could no longer be seen was calculated. This difference was added to the average shade to determine the final shade for the obstacle.

**Figure 2.**
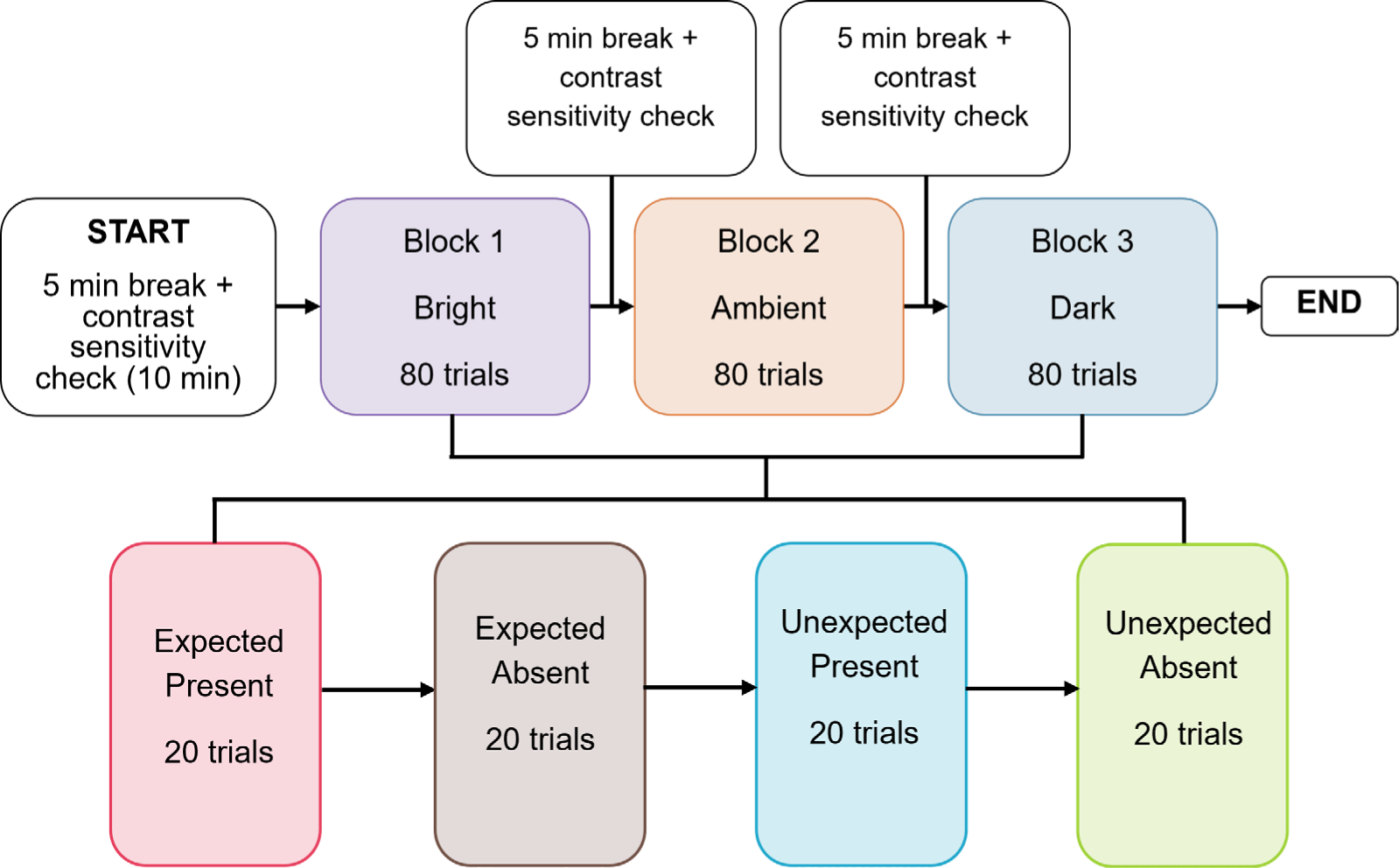
Illustration of the experiment structure.

The contrast sensitivity check took approximately 10 minutes and was preceded by the 5-minute break, providing a total of 15 minutes to pre-adapt to the light for each block. Additionally, it took approximately 25 minutes to complete one block, and the total duration of the experiment was no longer than 4 hours. Of this time, approximately 60 minutes were spent walking.

The obstacle presentation was controlled by streaming motion capture data from OptiTrack Motive (version 3.1) into Unity (version 2022). The obstacle was projected by a projector mounted on the ceiling. Lab Streaming Layer (LSL) was used to send triggers (or mark events) in the EEG data at two key points: before the participant stepped over the obstacle (the preparation phase) and after the obstacle was crossed (the reset phase). The preparation phase was defined as the window in which the participant left the starting area until they reached the obstacle. The reset phase was defined as the window in which the participant crossed the obstacle until they reached the end of the walkway. In trials where obstacles were absent, the preparation and reset phases were defined using a point in the centre of the walking path. The preparation phase corresponded to the distance between the start area and the centre point, and the reset phase to the distance between the centre point and the end area. The preparation and reset phases were used as temporal markers while analysing the EEG data.

### 2.3. EEG recording

EEG data were recorded using 32 Ag/AgCl electrodes which were connected to a portable amplifier (ANT Neuro, Hengelo, The Netherlands). Electrodes were positioned according to the International 10-20 system, and EEG signals were sampled at 500 Hz. The electrode impedances were reduced below 5 kΩ before starting the recording.

### 2.4. EEG preprocessing

EEG preprocessing was conducted using the MNE library (version 1.8.0) in Python (version 3.12). Data from the mastoid channels were removed, and a 1–100 Hz bandpass filter was applied to the remaining data. Linear trends were removed from the data to correct for slow drifts and improve signal quality. To identify channels containing prominent artifacts, those with kurtosis z-scores exceeding 5 were marked as bad and subsequently interpolated. On average, 0.6% of channels per participant were interpolated. Although CPz was used as the default reference while recording the data, the data were re-referenced to the average of all channels during preprocessing.

Independent Component Analysis (ICA) was performed using the extended infomax algorithm to identify and remove non-neural artifacts from the EEG data (Lee et al., 1999). Components were automatically classified using ICLabel (Li et al., 2022) which assigns probabilities for the following categories: brain, muscle artifact, eye blink, heartbeat, line noise, channel noise, and other. Components labelled as muscle with a probability ≥ 0.5 and any other non-brain components with a probability ≥ 0.7 were excluded. Additionally, all components classified as ‘other’ were excluded regardless of probability. Ocular artifacts were additionally identified by correlating ICA component time courses with frontal electrodes (FP1, FP2, F7 and F8) using MNE’s find_bads_eog function. A dynamic z-score threshold was applied starting at 3.5 and decreasing in steps of 0.05 until a minimum of 3 components were identified. Temporal autocorrelation was computed for each component using a 20 ms lag (Chaumon et al., 2015). Autocorrelation values were z-scored across components and components more than 1.5 SD below the mean were flagged as likely muscle artifacts and excluded. Focality of each component’s scalp topography was assessed using kurtosis (Nolan et al., 2010). Components with kurtosis z-scores exceeding 2 SD above the mean were excluded as overly focal and thus unlikely to reflect neural activity. Finally, manual inspection was conducted to validate or override automatic rejection where necessary. On average, 12 ± 3 (mean ± SD) non-artifactual components were retained for analysis. One participant was excluded at this stage due to excessive artifact contamination, _resulting_ in fewer than five non-artifactual components being retained after ICA. After artifact removal, the data were bandpass filtered between 1–40 Hz.

The EEG data were then segmented into 4 epochs: a 2 second standing baseline prior to walking onset, the preparation phase from walking onset to obstacle crossing onset (up to 9 seconds), the crossing period (up to 1.05 seconds), and a 1.3 second reset phase following the end of obstacle crossing. The first trial of each block was a practice trial and was removed. Trials missing key event markers, such as walking onset, obstacle crossing start or obstacle crossing end, were excluded. Additionally, trials lacking at least 1.3 seconds of data following obstacle crossing were discarded. To account for inter-individual variability, trials in which the crossing duration exceeded 1.05 seconds, or where the preparation phase exceeded 9 seconds, were also removed. Remaining epochs were cleaned using the AutoReject algorithm (Jas et al., 2017), which uses cross-validation to automatically identify and interpolate bad channels within epochs and rejects epochs that cannot be repaired. A maximum of 2 channels were permitted for interpolation per epoch. To address variability in the duration of the preparation phase, linear interpolation was used to time warp the preparation segment of each trial to match a global median duration specific to its condition (Supplementary Section S2, Table S1). This global median duration was derived by first computing each participant’s mean preparation duration for each condition, followed by calculating a global median across participants within each condition (Mustile et al., 2021). Following time warping, trials in which the preparation duration deviated excessively from the condition median were removed. Specifically, the ratio of the condition median duration to the trial’s actual preparation duration was computed for each trial, and trials where this ratio deviated more than ± 2.5 SD from the mean were excluded. The crossing segment was excluded from the final analysed epochs, which comprised the 2 second baseline, the time-warped preparation phase, and the 1.3 second reset phase. Remaining epochs were visually inspected, and those containing residual artifacts were removed. Participants who retained fewer than 50% of trials in any of the 12 Lighting × Obstacle conditions were excluded from analysis, resulting in the removal of one additional participant. On average, 18 ± 2 epochs per condition were retained for analysis across the remaining 42 participants, corresponding to an overall data loss of 10.7%. Event-Related Spectral Perturbations (ERSPs) were computed using Morlet wavelet Time-Frequency Representations (TFRs) across frequencies from 4–30 Hz. For each trial, a log-ratio baseline correction was applied relative to the 2 second standing period at the start of the trial. The baseline-corrected values were then averaged across trials and converted to percentage change from baseline (Supplementary Section S3, Figures S1 and S2).

### 2.5. EEG statistical analysis

Statistical differences in time-frequency power were assessed using non-parametric cluster-based permutation tests (Maris & Oostenveld, 2007) over channel × frequency × time space. Analysis covered 4–30 Hz, the last 2.7s of the preparation phase, and the first 1s of the reset phase. Electrode adjacency was defined using Delaunay triangulation and combined with frequency and time adjacency to form a three-dimensional adjacency matrix. Separate adjacency matrices were computed for the preparation and reset segments due to differences in their temporal extent. A cluster-forming threshold of α = 0.001 was applied at each channel-frequency-timepoint, and datapoints exceeding this threshold were grouped into clusters. For each cluster, a cluster-level statistic was calculated as the sum of test statistics (*t*_sum_ or *F*_sum_ cluster mass). A null distribution was then generated using 1000 permutations; on each permutation clusters were identified and only the maximum cluster mass was retained (Jas et al., 2018). The probability of each observed cluster was computed as the proportion of permutation-derived maximum cluster masses that were greater than the observed cluster mass, controlling the family-wise error rate (Maris & Oostenveld, 2007). Clusters with *p* < .05 were considered significant. Effect sizes were computed following Meyer et al. (2021), using a rectangular bounding box approach applied across the channel, frequency, and time dimensions of each significant cluster.

Statistical tests were performed for the following factors: Age (Young vs Older) was tested using an independent-samples t-test with a two-tailed threshold (permutation_cluster_test function). Under the null hypothesis, the data in the two age groups were drawn from the same probability distribution (Maris & Oostenveld, 2007). Lighting (3 levels: Bright, Ambient, Dark) and Obstacle (4 levels: Expected Present, Expected Absent, Unexpected Present, Unexpected Absent) were each tested using a one-way repeated-measures ANOVA (permutation_cluster_test with f_mway_rm) with a one-tailed threshold. For the Lighting analysis, each subject’s data were averaged across the four obstacle conditions. For the Obstacle analysis, each subject’s data were averaged across the three lighting conditions. Under the null hypothesis, the data in all levels of each factor were exchangeable (Maris & Oostenveld, 2007; Sassenhagen & Draschkow, 2019). Effect size for Age was quantified using Cohen’s *d*. Effect sizes for Lighting and Obstacle were quantified as partial eta-squared, recomputed by re-running the repeated measures ANOVA on data averaged within the rectangular cluster bounding box.

Conditional on a significant omnibus effect, post-hoc tests were conducted within a region of interest (ROI) defined by the electrodes and frequencies of the omnibus cluster across the full preparation and reset period. Within this ROI, paired differences between conditions were computed for each subject and passed to one-sample t-tests (permutation_cluster_1samp_test) using time-only adjacency and a two-tailed cluster-forming threshold of α = 0.001. Under the null hypothesis, the signs of the paired differences were randomly flipped on each permutation (Jas et al., 2018). Only post-hoc clusters falling within the omnibus cluster’s significant time window are reported. Post-hoc contrasts for Lighting were: Bright vs Dark, Bright vs Ambient, and Ambient vs Dark. Post-hoc contrasts for Obstacle were: Present vs Absent, Unexpected Present vs Expected Present, Unexpected Absent vs Expected Absent, Unexpected Present vs Absent, and Expected Present vs Absent. Present was defined as the average of Expected Present and Unexpected Present, and Absent was defined as the average of Unexpected Absent and Expected Absent.

Age × Lighting interactions were examined using targeted mixed ANOVAs during the preparation phase, comparing Bright and Dark conditions with predefined ROIs. Analyses focused on frontal (FP1, FPz, FP2, F7, F3, Fz, F4, F8) theta (4–7 Hz), occipito-parietal (P7, P3, Pz, P4, P8, POz, O1, Oz, O2) alpha (8–12 Hz), and fronto-central (FC5, FC1, FC2, FC6, C3, Cz, C4, CP5, CP1, CP2, CP6) beta (13–30 Hz) power. Targeted paired t-tests were conducted during the preparation phase to compare frontal theta (4–7 Hz) power between Bright and Dark conditions for Expected Present and Unexpected Present trials (see Supplementary Section S4).

### 2.6. Behavioural data processing and analysis

Motion capture data were recorded using the OptiTrack system (10 Prime × 13 cameras) and Motive software with a sampling rate of 240 Hz. Gait processing was performed in Python (version 3.11) using custom scripts. Trials were excluded if key timing events were missing (e.g., no recorded crossing event, missing walk onset), if events were not logged in the expected order of walk onset, crossing start, and crossing end, if the preparation time exceeded 9 s, or if the crossing duration exceeded 1.05 s. Following these exclusions, trials in which the preparation duration or crossing duration deviated by more than 2.5 SD from the participant’s condition mean were removed as outliers. Any participant who retained less than half of the trials in any condition after these exclusions was removed from the analysis. This resulted in the exclusion of one older adult participant. The final sample for gait analysis comprised 22 young adults and 19 older adults.

Motion capture data for the pelvis, left foot and right foot bone were extracted from each trial. Gaps in bone trajectories of up to 15 frames were filled using linear interpolation. The anterior-posterior position of each foot relative to the pelvis was computed and linearly detrended. Left and right heel strikes were identified as the positive peaks of the anterior-posterior position of the left or right foot bone relative to the pelvis bone. Detected heel strikes were visually inspected and manually corrected where necessary. Following correction, trials with less than 6 heel strikes (5 steps) were excluded, and it was confirmed that retained trials contained alternating left and right heel strikes.

Motion capture data were filtered using a zero-lag 4^th^ order low-pass Butterworth filter with a cutoff frequency of 6 Hz. Spatiotemporal gait parameters were computed for each trial as follows. Stride length was defined as the absolute anterior-posterior displacement of the foot between two consecutive heel strikes of the same foot. Stride time was defined as the temporal interval between two consecutive heel strikes of the same foot. Gait speed was computed as stride length divided by stride time. Cadence was defined as the number of steps (each step being the interval between consecutive heel strikes) divided by the walking duration, and expressed in steps per minute. Physiologically implausible values were used to identify outliers. Trials with cadence ≤ 60 or ≥ 141 steps per minute were excluded.

Individual strides with stride length ≤ 0.6 m or ≥ 1.7 m, stride time ≤ 0.85 s or ≥ 2 s, or speed ≤ 0.3 m/s or ≥ 2 m/s were removed, and any trials with fewer than 2 strides remaining were excluded.

Speed and cadence were computed from the same trials that contributed to EEG analysis. For each participant, all speed values within each Lighting × Obstacle condition were averaged, and cadence was averaged across all trials within each Lighting × Obstacle condition. A minimum of 32 strides per condition were retained for speed analysis, and a minimum of 10 trials per condition for cadence analysis.

Leg length was computed from the Motive skeleton model as the sum of 3 bone segment distances: pelvis-to-thigh, thigh-to-shin, and shin-to-foot. These were calculated separately for the left and right legs using the 3D Euclidean distance between bone positions and then averaged to obtain a single leg length estimate per participant. A Welch’s independent-samples t-test showed no between-group difference in leg length, *t*(27.81) = 0.18, *p* = .86, and there was no difference in height between young and older adults (*U* = 267.0, *p* = .13). As there were no between-group differences in either leg length or height, gait parameters were not normalised.

Gait speed and cadence were analysed with linear mixed-effects models initially fitted with the lme4 package (Bates et al., 2015) in R (version 4.2.2; R Core Team, 2022). Each model included the fixed effects of Age (Young, Older) and Lighting (Bright, Ambient, Dark), and their interaction, as well as Obstacle (Expected Present, Expected Absent, Unexpected Present, Unexpected Absent). Participants were also included as random intercepts. Model assumptions were checked by inspecting residuals for normality and homogeneity of variance. Residual variance heteroscedasticity was detected, and models were refitted using the nlme package (Pinheiro et al., 2022) to allow unequal residual variance. For speed allowing different residual variances per light condition significantly improved model fit (*p* < .001) and was retained as the final model. For cadence, allowing different residual variances per obstacle condition provided the best fit (*p* = .002) and was retained as the final model. Significant effects of Lighting were followed up with pairwise comparisons, and significant effects of Obstacle were followed up with contrasts matching the EEG post-hoc structure, using estimated marginal means (emmeans package; Lenth & Piaskowski, 2025) with Bonferroni correction.

## 3. Results

### 3.1. Walking speed and cadence

Means (standard errors) for gait speed and cadence are reported in Supplementary Section S5, Table S2.

Young adults walked faster than older adults (main effect of age, *F*(1,39) = 16.38, *p* < .001, 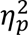 = .30). In contrast, cadence did not differ significantly between age groups (*F*(1,39) = 2.78, *p* = .104, 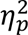 = .07). Lighting reliably affected gait: for both speed and cadence, participants moved more slowly in Dark lighting conditions. Speed showed a main effect of lighting (*F*(2,444) = 47.03, *p* < .001, 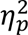 = .17), where walking was slower in the Dark than Bright, *t*(444) = 9.49, *p* < .001, and Ambient conditions, *t*(444) = 5.66, *p* < .001; walking was also faster in Bright than Ambient conditions, *t*(444) = 2.45, *p* = .044. Cadence showed a partially similar pattern (*F*(2,444) = 12.07, *p* < .001, 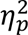 = .05), with cadence lower in the Dark than Bright, *t*(444) = 4.33, *p* < .001, and Ambient, *t*(444) = 3.94, *p* < .001, but no difference between Bright and Ambient, *t*(444) = 0.39, *p* = 1.0. No Age × Lighting interactions were observed for either speed (*F*(2,444) = 1.57, *p* = .209, 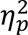= .01) or cadence (*F*(2,444) = 2.14, *p* = .119, 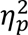 = .01).

Obstacle context also influenced both speed (*F*(3,444) = 32.77, *p* < .001, 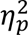 = .18) and cadence (*F*(3,444) = 35.57, *p* < .001, 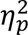 = .19). Participants walked faster when obstacles were Present compared to when they were Absent, *t*(444) = 4.60, *p* < .001, while cadence showed the opposite pattern, with higher cadence in Absent compared to Present trials, *t*(444) = 7.95, *p* < .001. Thus, an absence of obstacles was associated with shorter, more frequent steps (lower speed, higher cadence) compared to trials where obstacles were present.

Expectation further modulated gait. For speed, walking was faster for Expected Absent than Unexpected Absent trials, *t*(444) = 7.07, *p* < .001, and for Expected Present than Unexpected Present trials, *t*(444) = 5.21, *p* < .001. The same pattern was observed for cadence: Expected Absent was higher than Unexpected Absent, *t*(444) = 4.81, *p* < .001, and Expected Present higher than Unexpected Present, *t*(444) = 3.64, *p* = .002. Additionally, walking speed was faster for Expected Present than for Absent, *t*(444) = 6.76, *p* < .001, whereas Unexpected Present trials did not differ from Absent, *t*(444) = 0.74, *p* = 1.0. For cadence, both present conditions differed from Absent: Unexpected Present showed lower cadence than Absent, *_t_*(444) = 7.85, *p* < .001, and Expected Present was also lower than Absent, *t*(444) = 4.96, *p* < .001.

### 3.2. EEG results

#### 3.2.1. Preparation phase

##### Main effect of age

A negative cluster was observed (*t*_sum_ = −1,750.92, *p* = .039, *d* = −1.14), within the alpha band (12.07–12.97 Hz) over frontal and fronto-central electrodes (Fz, F4, FC1, FC2), approximately −2626 to −2312 ms before obstacle crossing (Figure 3). Baseline alpha power at these electrodes did not differ between young and older adults, Welch *t*(39) = 1.65, *p* = .108. Relative to baseline, alpha power was more desynchronised in the young group (−63.54%) compared to the older group (−48.43%).

**Figure 3.**
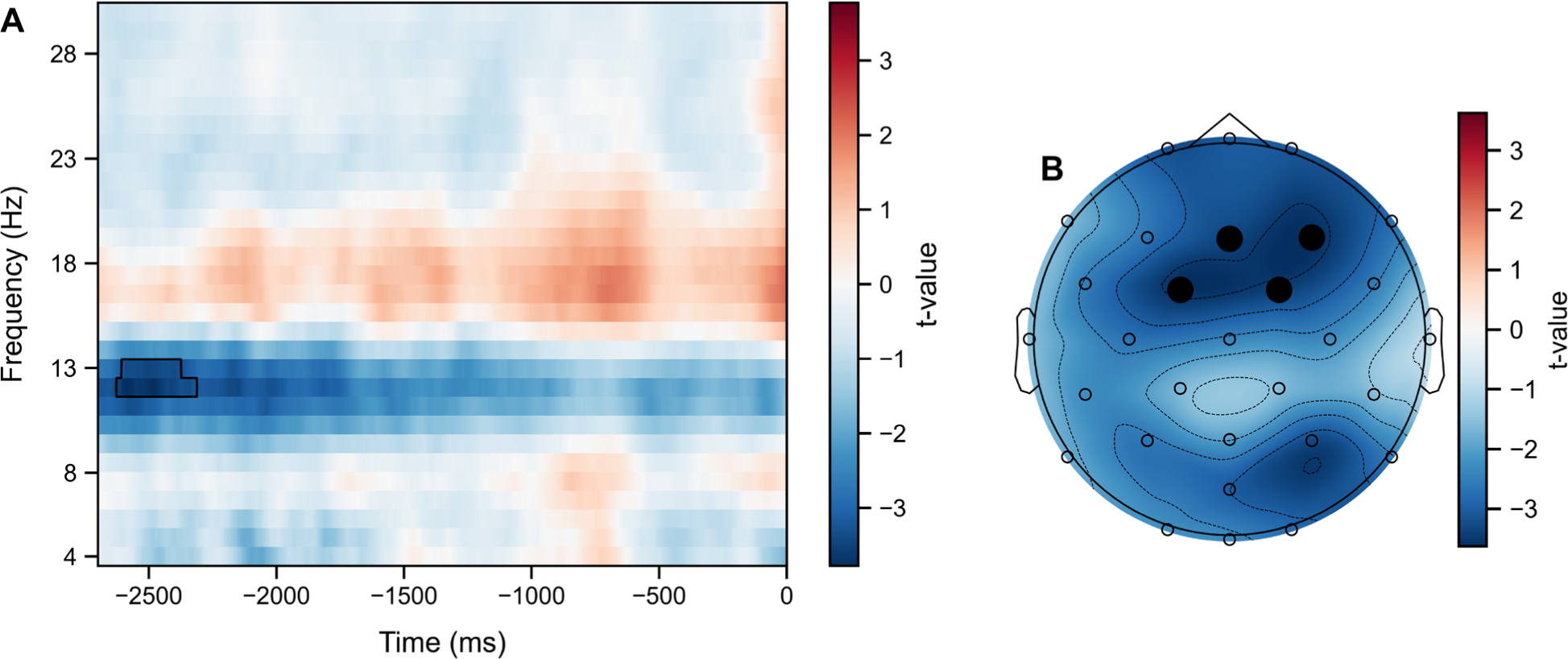
(A) Time-frequency representation of t-values averaged across significant cluster channels. The black contour delineates the significant cluster. Time 0 represents the start of obstacle crossing. (B) Topographic distribution of mean t-values across the cluster’s significant time-frequency window. Black dots indicate electrodes contributing to the cluster.

##### Main effect of lighting

Two clusters were observed during the preparation phase (Figure 4). The first cluster (*F*_sum_ = 434,950.30, *p* = .002, 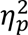 = .16) spanned the theta-alpha-beta frequency range (6.69–14.76 Hz), predominantly in the alpha band, across all electrodes (FP1, FPz, FP2, F7, F3, Fz, F4, F8, FC5, FC1, FC2, FC6, T7, C3, Cz, C4, T8, CP5, CP1, CP2, CP6, P7, P3, Pz, P4, P8, POz, O1, Oz, O2), approximately −2698 to −538 ms before obstacle crossing. Relative to baseline, power desynchronised in the Bright (−49.06%), Ambient (−54.03%), and Dark (−60.03%) conditions. Post-hoc comparisons identified that Ambient lighting induced more desynchronisation than Bright lighting from −1740 to −1676 ms (*t*_sum_ = 123.65, *p* = .013). Dark lighting elicited more desynchronisation than Bright across three windows: −2052 to −668 ms (*t*_sum_ = 3,237.44, *p* = .001), −2440 to −2140 ms (*t*_sum_ = 600.86, *p* = .002), and −2698 to −2498 ms (*t*_sum_ = 461.82, *p* = .002). Dark also showed more desynchronisation than Ambient from −2682 to −2598 ms (*t*_sum_ = 159.49, *p* = .006).

**Figure 4.**
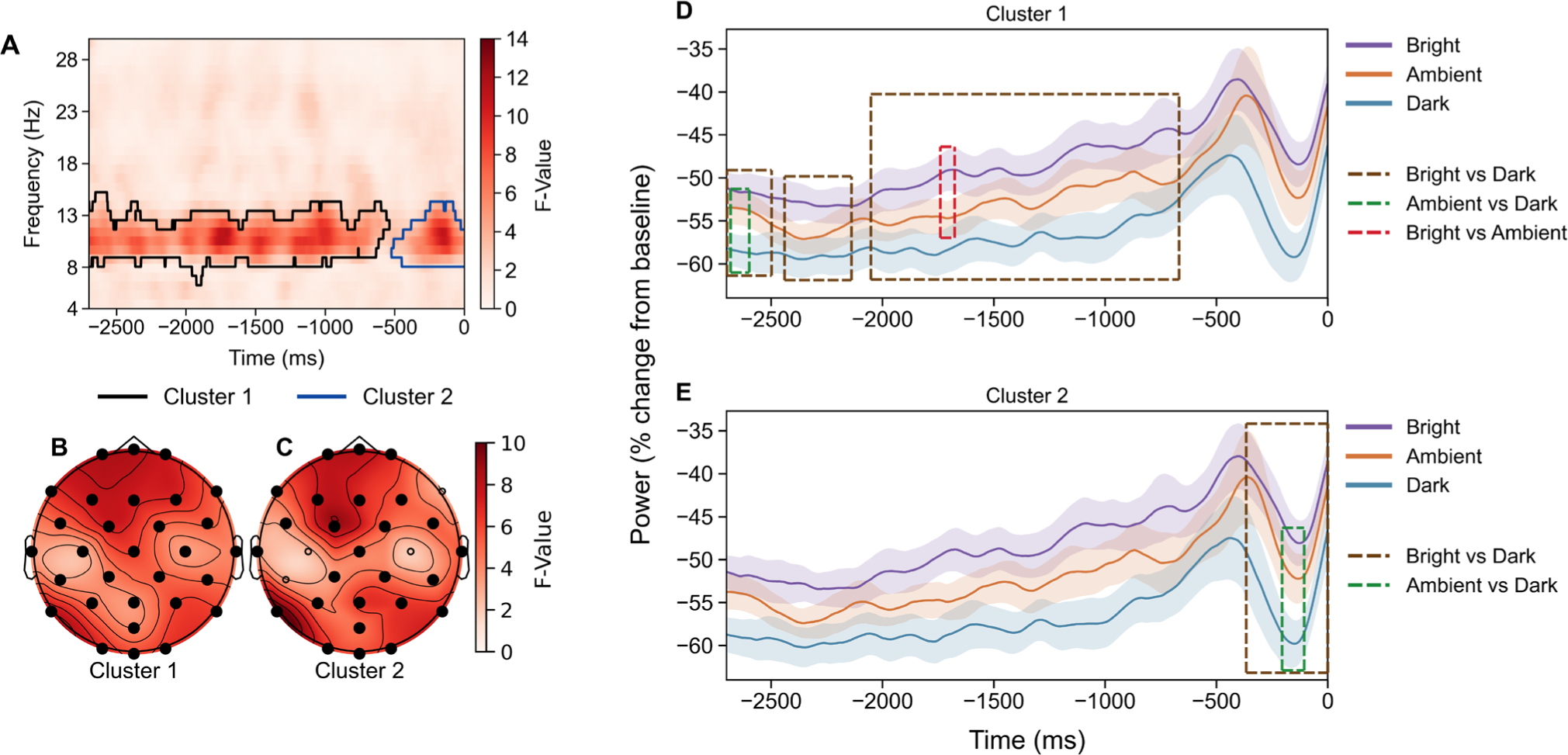
(A) Time-frequency representation of F-values averaged across all channels. Coloured outlines delineate the two significant clusters (Clusters 1 and 2), each shown in a distinct colour. Time 0 represents the start of obstacle crossing. (B–C) Topographic distributions of mean F-values across each cluster’s significant time-frequency window, for Cluster 1 and Cluster 2 respectively. Black dots indicate electrodes contributing to each cluster. (D–E) Mean power (percentage change from baseline) over time for each lighting condition, for (D) Cluster 1 and (E) Cluster 2. Lines show group means for Bright, Ambient, and Dark conditions. Shaded bands represent 95% within-subject confidence intervals (Cousineau-Morey method). Dashed boxes indicate time windows of significant post-hoc differences, colour-coded by contrast.

A second widespread cluster (*F*_sum_ = 101,711.11, *p* = .005, 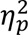= .18) was observed in the alpha-beta frequency range (8.48–13.86 Hz), predominantly in the alpha band, across frontal, temporal, central, parietal, and occipital electrodes (FP1, FPz, FP2, F7, F3, Fz, F4, FC5, FC1, FC2, FC6, T7, Cz, T8, CP1, CP2, CP6, P7, P3, Pz, P4, P8, POz, O1, Oz, O2), approximately −522 to 0 ms before crossing. Power desynchronised relative to baseline in the Bright (−45.47%), Ambient (−49.97%), and Dark (−58.79%) conditions.

Within Cluster 2, Dark lighting induced more desynchronisation than Bright from −366 to 0 _ms_ (*t*_sum_ = 933.31, *p* = .002), and Dark showed more desynchronisation than Ambient from −206 to −106 ms (*t*_sum_ = 188.93, *p* = .008). No significant time windows were observed for Bright vs Ambient conditions.

Of note, baseline alpha power differed across lighting conditions: Dark (22.07 dB) was higher than Bright (21.32 dB), *t*(41) = 3.51, *p* = .001, and higher than Ambient (21.41 dB), *t*(41) = 4.11, *p* < .001, but there was no difference between the Bright and Ambient conditions, *t*(41) = −0.63, *p* = .533.

##### Main effect of obstacle

Four clusters were observed during the preparation phase (Figure 5). The largest cluster (*F*_sum_ = 568,917.83, *p* = .003, 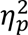 = .27) spanned the theta-alpha-beta range (4.00–30.00 Hz) across all electrodes (FP1, FPz, FP2, F7, F3, Fz, F4, F8, FC5, FC1, FC2, FC6, T7, C3, Cz, C4, T8, CP5, CP1, CP2, CP6, P7, P3, Pz, P4, P8, POz, O1, Oz, O2), approximately −1230 to −230 ms before obstacle crossing. Relative to baseline, power desynchronised in all four obstacle conditions: Expected Present (−27.58%), Expected Absent (−41.74%), Unexpected Present (−26.49%), and Unexpected Absent (−42.90%). Post-hoc comparisons for Cluster 1 revealed that Present trials showed less desynchronisation than Absent from −930 to −264 ms (*t*_sum_ = 1,502.95, *p* = .001). Unexpected Present and Expected Present did not differ significantly, nor did Unexpected Absent and Expected Absent. Unexpected Present also showed less desynchronisation than Absent from −922 to −274 ms (*t*_sum_ = 1,417.03, *p* = .001). Similarly, Expected Present showed less desynchronisation than Absent from −544 to −274 ms (*t*_sum_ = 609.35, *p* = .001) and from −882 to −738 ms (*t*_sum_ = 279.43, *p* = .004).

**Figure 5.**
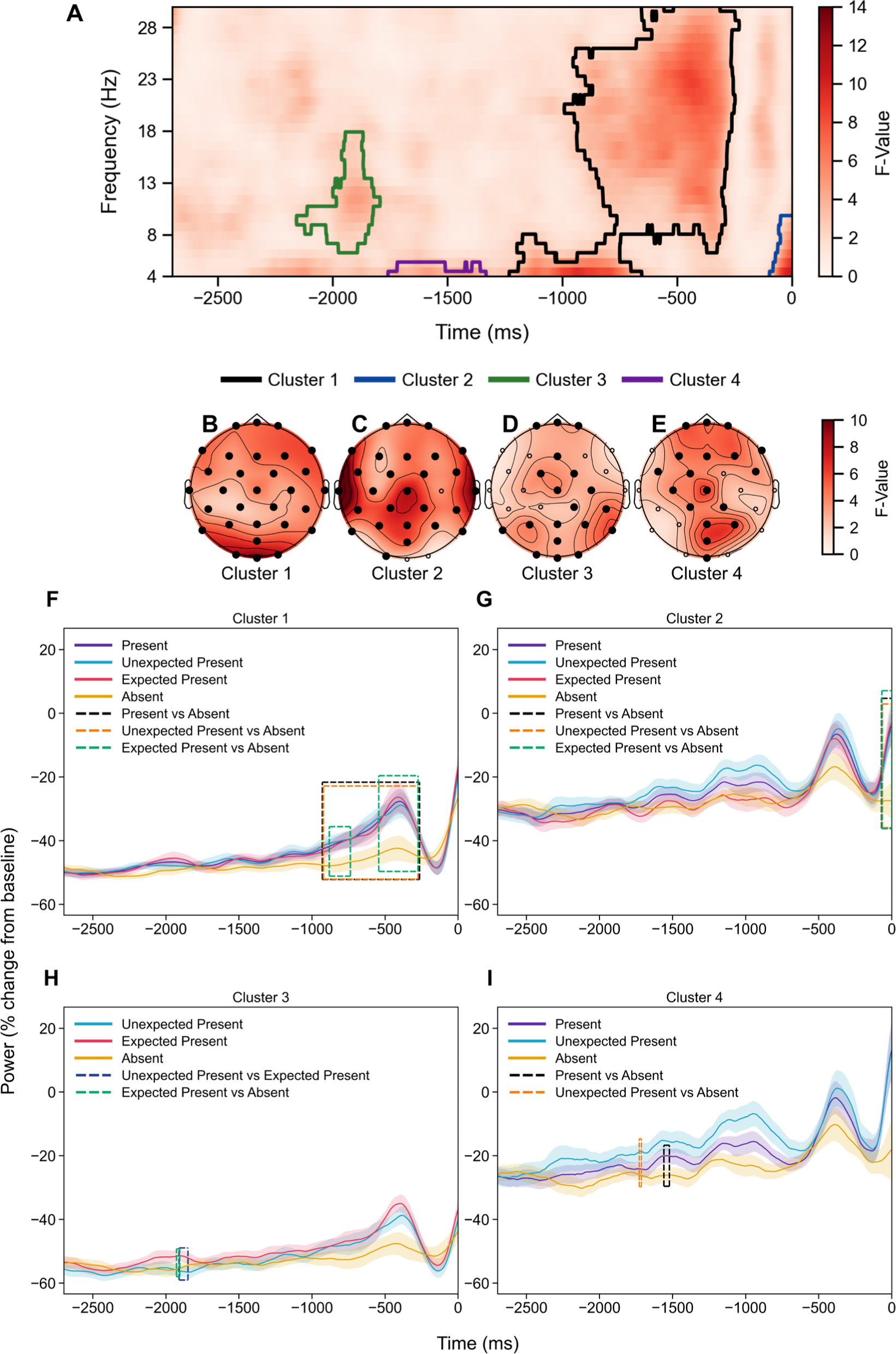
(A) Time-frequency representation of F-values averaged across all channels. Coloured outlines delineate the four significant clusters (Clusters 1–4), each shown in a distinct colour. Time 0 represents the start of obstacle crossing. (B–E) Topographic distributions of mean F-values across each cluster’s significant time-frequency window, for Clusters 1–4 respectively. Black dots indicate electrodes contributing to each cluster. (F–I) Mean power (percentage change from baseline) over time for significant post-hoc comparisons within each cluster, for (F) Cluster 1, (G) Cluster 2, (H) Cluster 3, and (I) Cluster 4. Lines show condition means; shaded bands represent 95% within-subject confidence intervals (Cousineau-Morey method). Dashed boxes indicate time windows of significant post-hoc differences, colour-coded by contrast.

A second cluster (*F*_sum_ = 22,115.76, *p* = .025, 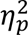 = .20) was identified in the theta-alpha range (4.00–9.38 Hz) across frontal, central, temporal, parietal, and occipital electrodes (FP1, FPz, FP2, F7, F3, Fz, F4, F8, FC5, FC1, FC2, FC6, T7, C3, Cz, T8, CP5, CP1, CP2, CP6, P7, P3, Pz, P4, P8, POz, O1), approximately −96 to 0 ms before crossing. Power desynchronised relative to baseline in Expected Present (−2.27%), Expected Absent (−25.70%), Unexpected Present (−4.19%), and Unexpected Absent (−24.80%) conditions. Within Cluster 2, Present trials exhibited less desynchronisation than Absent trials from −70 to 0 ms (*t*_sum_ = 189.92, *p* = .005). The Unexpected Present vs Expected Present and Unexpected Absent vs Expected Absent contrasts did not differ. Unexpected Present showed less desynchronisation than Absent from −64 to 0 ms (*t*_sum_ = 177.84, *p* = .003). Similarly, Expected Present trials had less desynchronisation than Absent trials between −68 to 0 ms (*t*_sum_ = 166.20, *p* = .006).

A third cluster (*F*_sum_ = 20,425.28, *p* = .029, 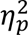 = .19) spanned the theta-alpha-beta range (6.69–17.45 Hz), predominantly in the alpha and beta bands, across frontal, central, parietal, and occipital electrodes (FP1, FPz, FP2, Fz, FC1, FC2, Cz, C4, CP2, CP6, P7, P3, Pz, P4, P8, POz, O1, Oz, O2), approximately −2156 to −1794 ms before crossing. Power desynchronised relative to baseline in Expected Present (−52.39%), Expected Absent (−52.58%), Unexpected Present (−57.47%), and Unexpected Absent (−60.11%) trials. No clusters were observed for Present vs Absent or Unexpected Absent vs Expected Absent within Cluster 3. When obstacles appeared unexpectedly (Unexpected Present), power was more desynchronised than when obstacles were expected (Expected Present), from −1910 to −1850 ms (*t*_sum_ = −115.50, *p* = .009). Unexpected Present and Absent showed no significant difference. Expected Present exhibited less desynchronisation than Absent from −1928 to −1906 ms (*t*_sum_ = 42.97, *p* = .016).

The fourth cluster (*F*_sum_ = 10,927.55, *p* = .045, 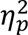 = .27) was restricted to the theta band (4.00–4.90 Hz) across frontal, central, parietal, and occipital electrodes (FP1, FPz, FP2, F3, Fz, F4, F8, FC5, FC1, FC6, C3, Cz, CP1, CP2, Pz, P4, POz, Oz), approximately −1758 to −1332 ms before crossing. Relative to baseline, power desynchronised in Expected Present (−28.06%), Expected Absent (−28.22%), Unexpected Present (−15.51%), and Unexpected Absent (−28.08%) conditions. Within Cluster 4, Present showed less desynchronisation than Absent from −1560 to −1522 ms (*t*_sum_ = 72.22, *p* = .015). No significant differences were found for Unexpected Present vs Expected Present or Unexpected Absent vs Expected Absent. When obstacles appeared unexpectedly (Unexpected Present), power was less desynchronised than in Absent trials, between −1728 to −1714 ms (*t*_sum_ = 28.55, *p* = .024). No significant time windows were identified between Expected Present and Absent trials.

#### 3.2.2. Reset phase

##### Main effect of age

No clusters were observed for the main effect of age during the reset phase.

##### Main effect of lighting

Two clusters were identified during the reset phase (Figure 6). The first (*F*_sum_ = 33,990.70, *p* = .007, 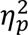 = .23) was observed in the alpha-beta frequency range (7.59–13.86 Hz), predominantly in the alpha band, across parietal and occipital electrodes (Pz, P4, P8, POz, O1, Oz, O2), approximately 456–998 ms after obstacle crossing. Relative to baseline, power desynchronised in the Bright (−29.75%), Ambient (−32.47%), and Dark (−45.25%) conditions. Post-hoc comparisons for the first cluster identified that Dark lighting induced more desynchronisation than Bright from 466 to 948 ms (*t*_sum_ = 1,182.17, *p* = .001), and Dark produced more desynchronisation than Ambient from 662 to 860 ms (*t*_sum_ = 402.10, *p* = .002). There were no significant differences between Bright and Ambient trials.

**Figure 6.**
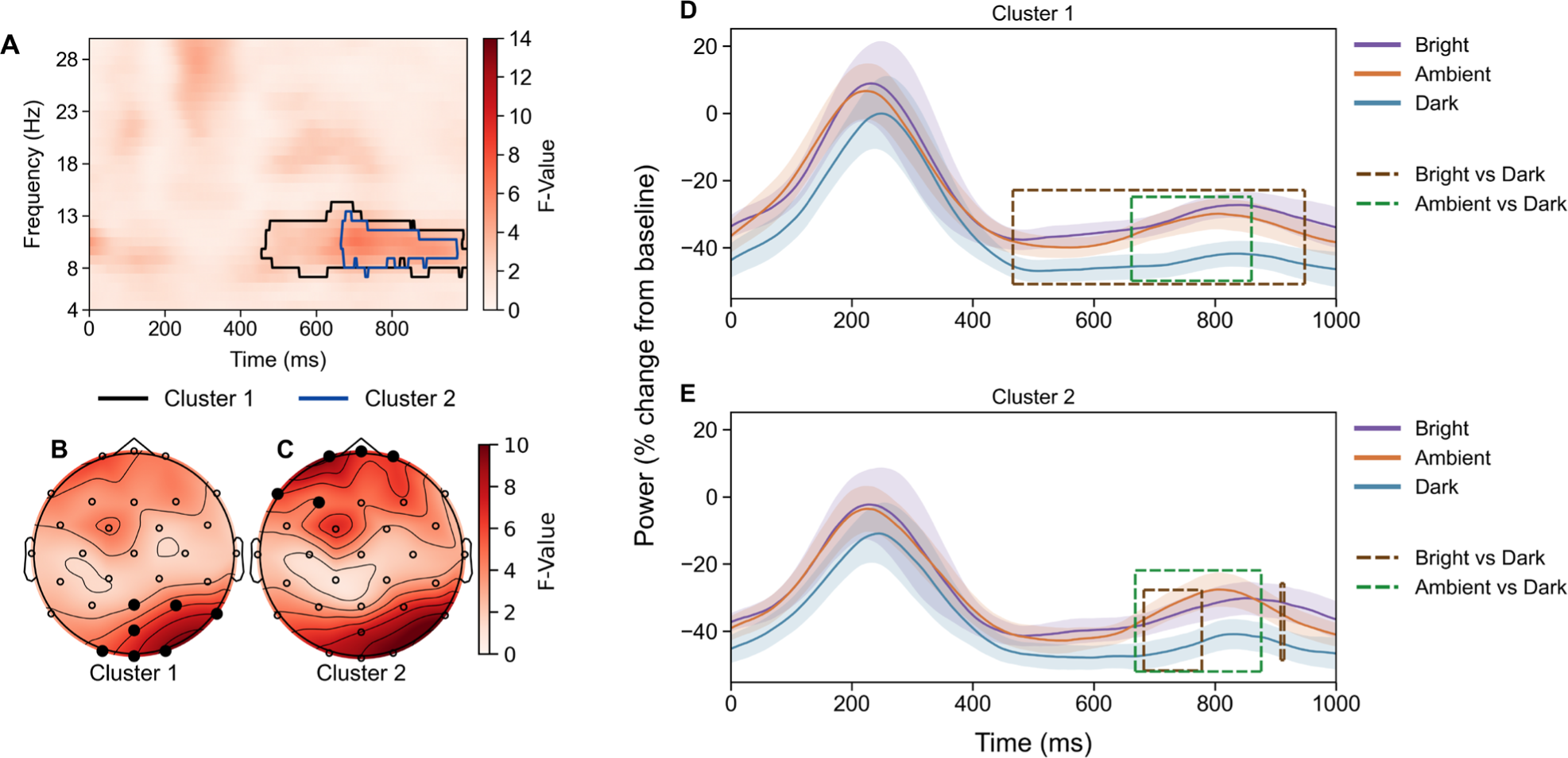
(A) Time-frequency representation of F-values averaged across all channels. Coloured outlines delineate the two significant clusters (Cluster 1 and Cluster 2), each shown in a distinct colour. Time 0 represents the end of obstacle crossing. (B–C) Topographic distributions of mean F-values across each cluster’s significant time-frequency window, for Cluster 1 and Cluster 2 respectively. Black dots indicate electrodes contributing to each cluster. (D–E) Mean power (percentage change from baseline) over time for each lighting condition, for (D) Cluster 1 and (E) Cluster 2. Lines show group means for Bright, Ambient, and Dark conditions. Shaded bands represent 95% within-subject confidence intervals (Cousineau-Morey method). Dashed boxes indicate time windows of significant post-hoc differences, colour-coded by contrast.

A second cluster (*F*_sum_ = 8,380.19, *p* = .023, 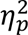 = .18) was also observed within the alpha band (7.59–12.97 Hz) across frontal electrodes (FP1, FPz, FP2, F7, F3), approximately 666– 972 ms after crossing. Power desynchronised relative to baseline in the Bright (−32.55%), Ambient (−31.99%), and Dark (−45.68%) conditions. Within Cluster 2, Dark lighting showed more desynchronisation than Bright, between 682 to 778 ms (*t*_sum_ = 181.00, *p* = .001) and from 908 to 914 ms (*t*_sum_ = 14.20, *p* = .008), and Dark induced more desynchronisation than Ambient from 668 to 876 ms (*t*_sum_ = 435.61, *p* = .001). No significant differences were observed between Bright and Ambient lighting.

##### Main effect of obstacle

A single cluster (Figure 7) was observed (*F*_sum_ = 2,783,777.89, *p* = .001, 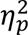 = .54), spanning the theta-alpha-beta range (4.00–30.00 Hz) across all electrodes (FP1, FPz, FP2, F7, F3, Fz, F4, F8, FC5, FC1, FC2, FC6, T7, C3, Cz, C4, T8, CP5, CP1, CP2, CP6, P7, P3, Pz, P4, P8, POz, O1, Oz, O2), from 0 to 998 ms after obstacle crossing. Relative to baseline, power synchronised in Expected Present (23.14%) and Unexpected Present (26.29%) trials and desynchronised in Expected Absent (−26.42%) and Unexpected Absent (−23.20%) trials.

**Figure 7.**
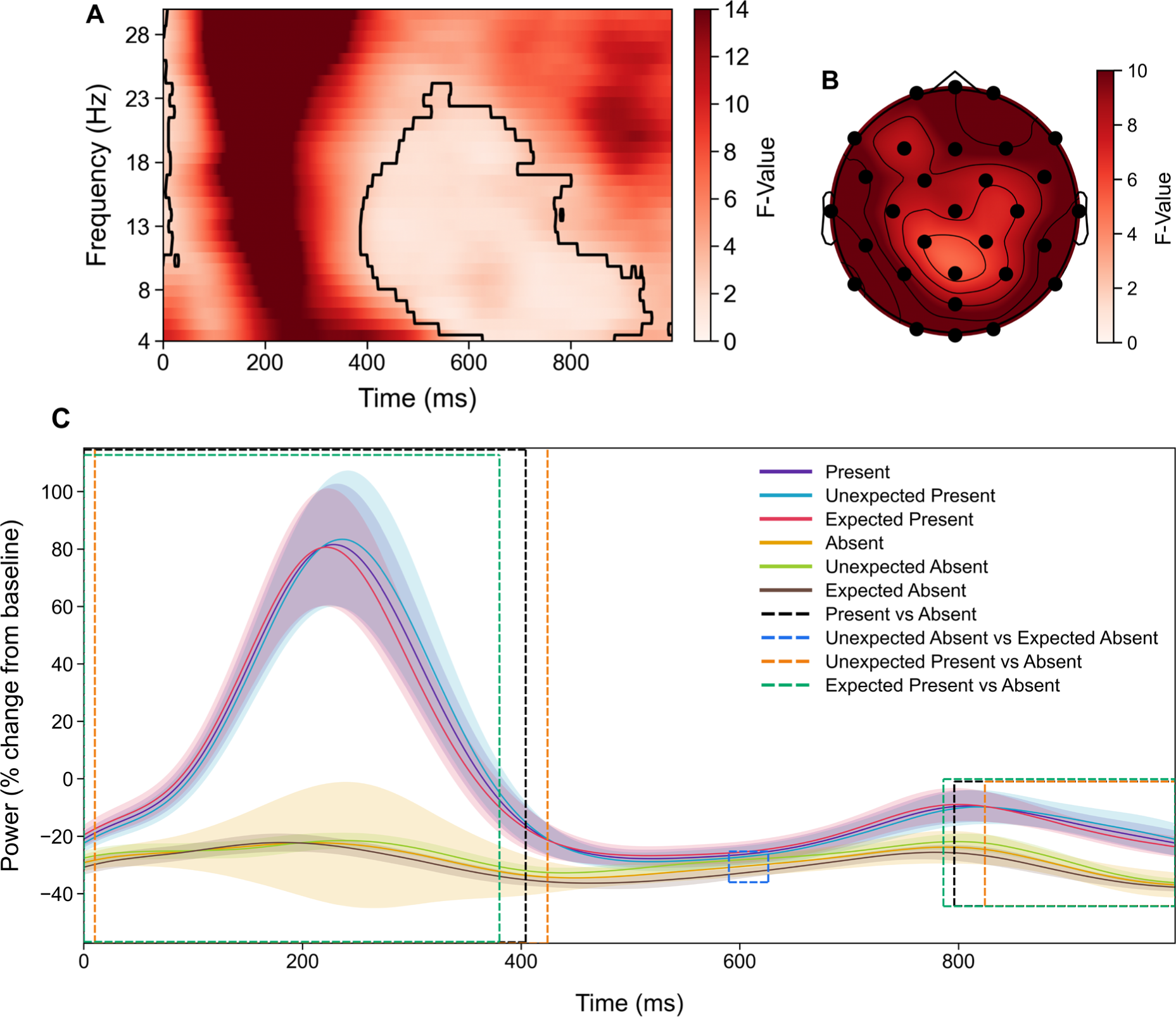
(A) Time-frequency representation of F-values averaged across all channels. The black contour delineates the significant cluster. Time 0 represents the end of obstacle crossing. (B) Topographic distribution of mean F-values across the cluster’s significant time-frequency window. Black dots indicate electrodes contributing to the cluster. (C) Mean power (percentage change from baseline) over time for post-hoc comparisons within the obstacle cluster (Present vs Absent, Unexpected Absent vs Expected Absent, Unexpected Present vs Absent, and Expected Present vs Absent). Shaded bands represent 95% within-subject confidence intervals (Cousineau-Morey method). Dashed boxes indicate time windows of significant post-hoc differences, colour-coded by contrast.

Post-hoc comparisons revealed that following obstacle crossing, power synchronised in Present and desynchronised in Absent trials between 0 to 404 ms (*t*_sum_ = 1,190.46, *p* = .001). There was less desynchronisation in Present trials relative to Absent, between 796 to 998 ms (*t*_sum_ = 499.90, *p* = .001). Unexpected Present and Expected Present did not differ. Unexpected Absent trials had less desynchronisation than Expected Absent from 590 to 626 ms (*t*_sum_ = 69.37, *p* = .006). When obstacles appeared unexpectedly (Unexpected Present), power synchronised while power desynchronised in Absent trials from 10 to 424 ms (*t*_sum_ = 1,108.34, *p* = .001). Unexpected Present trials also induced less desynchronisation than Absent trials from 824 to 998 ms (*t*_sum_ = 393.05, *p* = .001). Power synchronised when obstacles were expected (Expected Present) and desynchronised in Absent trials from 0 to 380 ms (*t*_sum_ = 1,148.16, *p* = .001). Expected Present trials also showed less desynchronisation than Absent trials from 786 to 998 ms (*t*_sum_ = 544.38, *p* = .001).

## 4. Discussion

We investigated the effects of 3 different lighting conditions (Bright, Ambient and Dark) on the neural and behavioural dynamics of walking and obstacle crossing in young and older adults. As expected, we identified that reduced visibility led to frequency-specific changes in cortical activity during both the crossing preparation and reset phases. Dark conditions reliably induced alpha desynchronisation, indicating increased engagement of the visual and attention networks when visual input was reduced. The presence of obstacles induced broadband changes, with reduced desynchronisation across theta-alpha-beta bands during the crossing preparation phase, and a transient post-crossing synchronisation consistent with a motor resetting response. In contrast, and contrary to our predictions, age and expectation effects were small. Young adults had a slightly increased frontal alpha desynchronisation relative to older adults during the crossing preparation phase, and there was a short-lived and broad-band expectation effect when obstacles appeared unexpectedly, compared to when they were expected.

### 4.1. Reduced visibility increases attention demands during walking

The present findings show that reduced visibility substantially increased cortical engagement during walking, reflected by robust alpha desynchronisation across both preparation and reset phases. This partly aligns with the broader literature on how the brain adapts to degraded visual input during walking. For example, Cao et al. (2020) reported that occipital alpha power decreased during walking relative to standing regardless of whether participants walked in light or complete darkness, with alpha power being lower in the light than in the dark condition. The authors interpreted this reduction as a release of inhibition over peripheral visual processing, reflecting a shift in attentional state during walking. However, in the present study, Dark conditions elicited more alpha desynchronisation despite participants walking more slowly. Considering that alpha desynchronisation has been associated with cortical activation (Neuper & Pfurtscheller, 2001; Pfurtscheller, 2001), with greater desynchronisation reflecting engagement of a larger cortical network (Pfurtscheller & Lopes da Silva, 1999), this suggests that the alpha modulation observed here does not simply reflect the processing of visual input, but the increased cortical demand required when visual information is not sufficient to allow for proactive gait adjustments.

This interpretation is consistent with studies showing that when visual input is restricted, there is a shift towards sensory reweighting, increasing reliance on somatosensory information. Oliveira et al. (2017) found that walking with eyes closed produced stronger theta to beta desynchronisation over the bilateral somatosensory cortex, reflecting greater engagement of this area to maintain balance and co-ordinate gait. Similar patterns have been observed in tasks that challenge stability, such as balance beam walking, where beta power decreases over sensorimotor areas (Sipp et al., 2013), consistent with the idea that reduced beta reflects increased motor control demands (Liu et al., 2025). The present findings extend this by showing that even partial reductions in visual input, rather than complete occlusion, are enough to increase cortical engagement during motor preparation.

The strong alpha desynchronisation observed in Dark trials cannot be attributed entirely to walking speed. Faster walking typically produces greater desynchronisation across the alpha and beta bands (Nordin et al., 2020; Salminen et al., 2025), yet participants walked slowest in the Dark condition. The fact that desynchronisation increased despite slower gait strongly suggests that this is reflective of task-specific visuomotor demands, where cortical resources are increased even when behavioural adjustments (slower speed and reduced cadence) might otherwise reduce the overall cognitive load.

These findings indicate that reduced visibility imposes two challenges during walking. Dark conditions limit the availability of visual information needed for proactive adjustments, while simultaneously increasing the reliance on somatosensory and attention networks to maintain stability. This could explain why we observed widespread alpha desynchronisation under low lighting, whereas studies using complete occlusion or task-free walking have reported different patterns. The neural response to reduced visibility is therefore not a simple product of visual input but an interaction between sensory uncertainty and the anticipatory demands of obstacle negotiation.

### 4.2. Obstacle preparation and reset dynamics

In our previous work, we demonstrated that unexpected obstacles induced an increase in frontal theta power at the time of the obstacles appearing, reflecting rapid proactive control when participants have limited time to adjust their gait (Mustile et al., 2021). This was followed by a decrease in beta power over sensorimotor regions when participants approached the obstacle, interpreted as motor preparation, and a post-crossing beta rebound reflecting reactive resetting of the motor system.

In the current study, the presence of obstacles during the preparation phase produced a broadband reduction in desynchronisation across the theta-alpha-beta range relative to unobstructed walking. This aligns with Nordin et al. (2019), who reported increases in 3–13 Hz power over supplementary motor and premotor cortices shortly after obstacle appearance, followed by posterior parietal increases prior to crossing. These findings indicate that preparing to cross an obstacle engages a distributed cortical network that supports visuomotor planning and gait adaptation, even when obstacles are expected.

We found that unexpected obstacles induced greater desynchronisation in the alpha-beta range during preparation compared to expected obstacles. Greater alpha and beta desynchronisation reflects engagement of a larger cortical network (Pfurtscheller & Lopes da Silva, 1999). Therefore, these patterns suggest that when participants cannot predict whether or not an obstacle will appear, additional cortical resources are engaged to support proactive control. Notably, the expectation effect was small and transient. This highlights that heightened proactive control under uncertainty is dynamic and is easily overshadowed by the demands of obstacle negotiation itself.

The broadband nature of the preparation phase clusters highlights a methodological difference between our previous and present studies. In Mustile et al. (2021) we analysed spectral power within predefined frequency bands and cortical regions of interest and showed that theta and beta can show dissociated patterns during obstacle negotiation, with theta increasing at the time of the obstacle appearance and beta decreasing as participants reach the obstacle. In contrast, we used a cluster-based permutation approach in the present study which showed overall relatively higher power for obstacle-present trials, it is possible that opposite changes in adjacent narrow frequency bands (e.g., increased theta alongside decreased beta) were masked within the cluster approach.

During the reset phase, obstacle-present trials showed an early synchronisation above baseline followed by a gradual return towards desynchronisation. Although the cluster spanned the theta-alpha-beta range, the synchronisation observed in obstacle-present trials after crossing likely reflects the post-movement beta synchronisation or beta rebound (Pfurtscheller & Solis-Escalante, 2009; Pfurtscheller et al., 1996; Solis-Escalante et al., 2012). The beta rebound has been reported across a range of tasks, typically over sensorimotor regions, including voluntary hand movements (Pfurtscheller et al., 1998), balance perturbation (Nakamura et al., 2021), and postural sway during quiet stance (Nakamura et al., 2023). Although post-movement beta synchronisation has primarily been reported over the sensorimotor cortex (Pfurtscheller et al., 1996; Salmelin et al., 1995), involvement of pre- and post-central gyrus, supplementary motor area, and frontal medial cortex has also been observed (Parkes et al., 2006).

The beta rebound over sensorimotor areas has been proposed to reflect a recalibration of the motor system in preparation for the next movement (Kilavik et al., 2013). Additionally, the beta rebound has been linked to reactive motor control. Liebrand et al. (2017) reported increased beta over prefrontal and sensorimotor regions during reactive motor inhibition in a go/nogo task. In our previous work we extended this reactive control interpretation to obstacle navigation, observing a post obstacle crossing beta rebound over frontal, central, and parietal electrodes, with the strongest effects observed over parietal regions, which we interpreted as reflecting a reactive control mechanism resetting the motor system to its previous state (Mustile et al., 2021). The present findings are consistent with this interpretation, suggesting that the beta rebound observed after obstacle crossing reflects a resetting of the motor system following the gait adjustment required to step over the obstacle. The scalp-wide distribution of the present cluster is consistent with previous findings indicating the beta rebound is not restricted to sensorimotor regions (Parkes et al., 2006), and with the frontal, central, and parietal distribution we previously observed during obstacle navigation (Mustile et al., 2021). The later portion of the reset phase cluster where obstacle-present trials showed less desynchronisation than obstacle-absent trials could reflect the gradual re-establishment of control following the initial resetting response. Additionally, Unexpected Absent trials induced less desynchronisation than the Expected Absent trials, which is contrary to expectation, as both conditions involved unobstructed walking and no difference was anticipated in the reset phase.

Finally, the relatively higher theta power for obstacle-present compared to absent trials across both preparation and reset phases supports the interpretation that obstacle negotiation engages processes over and above gait modulation. Theta increases have been consistently linked to balance related demands during walking, including balance beam walking (Sipp et al., 2013), perturbations in balance (Peterson & Ferris, 2018) and walking on uneven ground (Liu et al., 2024). The relatively higher theta power for obstacle-present trials in our study likely reflects the integration of balance with visuomotor planning, particularly given that stepping over an obstacle introduces greater stability demands than unobstructed walking.

### 4.3. Age differences are present but not dominant

Although we expected that older adults would be particularly affected by low lighting conditions, the age effects observed in this study were modest, and smaller than the main effects of lighting and obstacle presence. During the preparation phase, young adults had greater alpha desynchronisation over frontal and fronto-central electrodes than older adults. Alpha desynchronisation is considered to reflect cortical activation associated with sensory, cognitive, and motor processing, with greater desynchronisation indicating recruitment of a larger cortical network (Pfurtscheller & Lopes da Silva, 1999). The greater frontal alpha desynchronisation in the younger group may therefore indicate stronger engagement of these networks during obstacle preparation.

However, this interpretation is complicated by the fact that young adults walked faster than older adults, and walking speed is an established modulator of cortical activity. Faster gait is associated with greater alpha and beta desynchronisation in the sensorimotor cortex during treadmill walking (Nordin et al., 2020; Salminen et al., 2025), as well as in posterior parietal and cuneus regions (Salminen et al., 2025). The absence of baseline alpha differences between age groups suggests that the effect is not attributable to baseline spectral differences, but the influence of speed remains a plausible explanation.

No age differences emerged during the reset phase, further indicating that age was not a primary determinant of the reactive control processes engaged after obstacle crossing. This pattern aligns with recent work showing that older adults may rely more heavily on cortical resources at higher walking speeds (Salminen et al., 2025) but that age differences are reduced when gait demands are slower and self-determined. Future work to match gait speed across age groups would address whether age-related differences in frontal alpha persist when speed differences are controlled.

### 4.4. Trial-mean vs standing baselines

The time course of obstacle-related effects in the present study differed from previous mobile EEG studies of obstacle navigation, and these may be largely attributed to the choice of baseline. In this study, we computed event-related spectral perturbations relative to a standing baseline, expressing spectral power during preparation and reset as a percentage change from a 2-second stationary period. In contrast, Nordin et al. (2019) and Mustile et al. (2021) used trial-mean baselines, subtracting the average log spectrum across the entire epoch. Trial-mean baselines remove activity that is sustained throughout the trial and therefore isolate transient deviations linked to specific events, such as obstacle appearance.

Using a standing baseline preserves both sustained and transient components of walking activity. Because walking itself produces alpha desynchronisation (Cao et al., 2020; Delval et al., 2020), both obstacle-present and obstacle-absent conditions showed desynchronisation throughout the preparation phase. Condition differences therefore appeared primarily as differences in the magnitude of desynchronisation, rather than as increases above baseline. This contrasts with the more phasic responses reported in Nordin et al. (2019) of power increases in the 3–13 Hz range within 200 ms of obstacle appearance, and with Mustile et al. (2021) who reported frontal theta increases specifically when the unexpected obstacle appeared on the path.

### 4.5. Limitations

Several limitations should be considered when interpreting the findings of the present study. First, although cluster permutation tests effectively control for multiple comparisons, they provide limited temporal precision (Sassenhagen & Draschkow, 2019). The time windows reported here should therefore be interpreted descriptively rather than as precise onsets.

Second, walking speed differed between age groups and across lighting and obstacle conditions. Self-selected speed was used to preserve ecological validity, but this introduced potential confounds, as speed modulates theta, alpha and beta power in different ways. Although several effects ran counter to what speed alone would predict, a contribution of gait cannot be ruled out. Future studies could match speed across conditions to isolate its contribution.

Third, the use of a 32-channel mobile EEG system limited spatial resolution and the number of independent components that could be extracted during the ICA cleaning stage (Klug & Gramann, 2021). As a result, some neural activity may have been merged with broader components, contributing to the broadband patterns observed in the cluster analysis. Higher density systems would allow for more reliable component separation and improve the interpretability of our observed effects.

Finally, the obstacles were projected onto the floor rather than physical obstacles. Although this allowed for more precise control over the timing and location, projected obstacles may not fully replicate the biomechanical demands of stepping over physical obstacles. Future work could extend the paradigm to include physical obstacles to assess whether similar dynamics emerge.

### 4.6. Conclusion

This study identified that the neural demands of walking are shaped more by reduced visibility and the need to negotiate obstacles rather than age alone. Across both preparation and reset phases, dim lighting elicited predominantly alpha desynchronisation, while obstacle presence produced broadband modulation of theta, alpha and beta activity, indicating that additional cortical resources are recruited when visual information is limited and when gait must be adapted. Expectation effect was present but modest, suggesting that uncertainty recruits additional cortical resources during preparation, but this effect was overshadowed by the physical demands of obstacle negotiation. Age-related differences were detectable only in frontal alpha during preparation indicating that ageing effects on cortical engagement are subtle in healthy adults under self-selected speed conditions. We show that neural control of walking is highly sensitive to environmental conditions and provide a foundation for future work to examine how these neural responses may differ in individuals at an elevated risk of falls, and how environmental interventions may support safer mobility in older age.

## 5. Data and code availability

The mobile EEG data (Kaul et al., 2026c, https://doi.org/10.5281/zenodo.22705594) and motion capture data (Kaul et al., 2026d, https://doi.org/10.5281/zenodo.22707865) are openly available on Zenodo. The EEG and behavioural analysis code (Kaul et al., 2026a, https://doi.org/10.5281/zenodo.22734641), the gait processing code (Kaul et al., 2026b, https://doi.org/10.5281/zenodo.22714504), and the obstacle brightness calibration task (Kaul et al., 2026e, https://doi.org/10.5281/zenodo.22714522) are archived on Zenodo from GitHub.

## 6. Author contributions

Danishta Kaul: Conceptualization, Methodology, Software, Formal Analysis, Investigation, Data Curation, Writing - Original Draft, Writing - Review & Editing, Visualization, Project Administration, Funding Acquisition. Magda Mustile: Conceptualization, Writing - Review & Editing. Mark Donoghue: Methodology, Software, Writing - Review & Editing. Alexander Brownlee: Conceptualization, Methodology, Writing - Review & Editing, Supervision, Funding Acquisition. Magdalena Ietswaart: Conceptualization, Methodology, Writing - Review & Editing, Supervision, Funding Acquisition. Gemma Learmonth: Conceptualization, Methodology, Writing - Review & Editing, Supervision, Funding Acquisition.

## 9. Acknowledgements

We would like to thank Dr Liivia-Mari Lember for assistance with EEG cap application during data collection, and Professor Paul Hibbard and Dr Ross Goutcher for their contributions to the design of the procedure used to determine the obstacle brightness threshold. We would also like to thank all participants who took part in the study.

