## Supplementary Materials for "Environmental Demands Outweigh Age Effects in Cortical Dynamics During Walking"

#### S1. Obstacle width calculation

Participant stride length was estimated from height using the equation: height (inches) × 0.413. The estimated stride length was then divided by two to calculate step length, which was used to determine obstacle width.

#### S2. Median preparation duration

**Table S1**

*Median preparation-phase duration (ms) for each condition.*

| **Condition** | **Preparation Phase Duration (ms)** |
|---|---|
| Bright Expected Present | 2960 |
| Bright Expected Absent | 2760 |
| Bright Unexpected Present | 3340 |
| Bright Unexpected Absent | 2860 |
| Ambient Expected Present | 2950 |
| Ambient Expected Absent | 2780 |
| Ambient Unexpected Present | 3060 |
| Ambient Unexpected Absent | 3020 |
| Dark Expected Present | 3170 |
| Dark Expected Absent | 3040 |
| Dark Unexpected Present | 3410 |
| Dark Unexpected Absent | 3380 |

### S3. Time-frequency representations

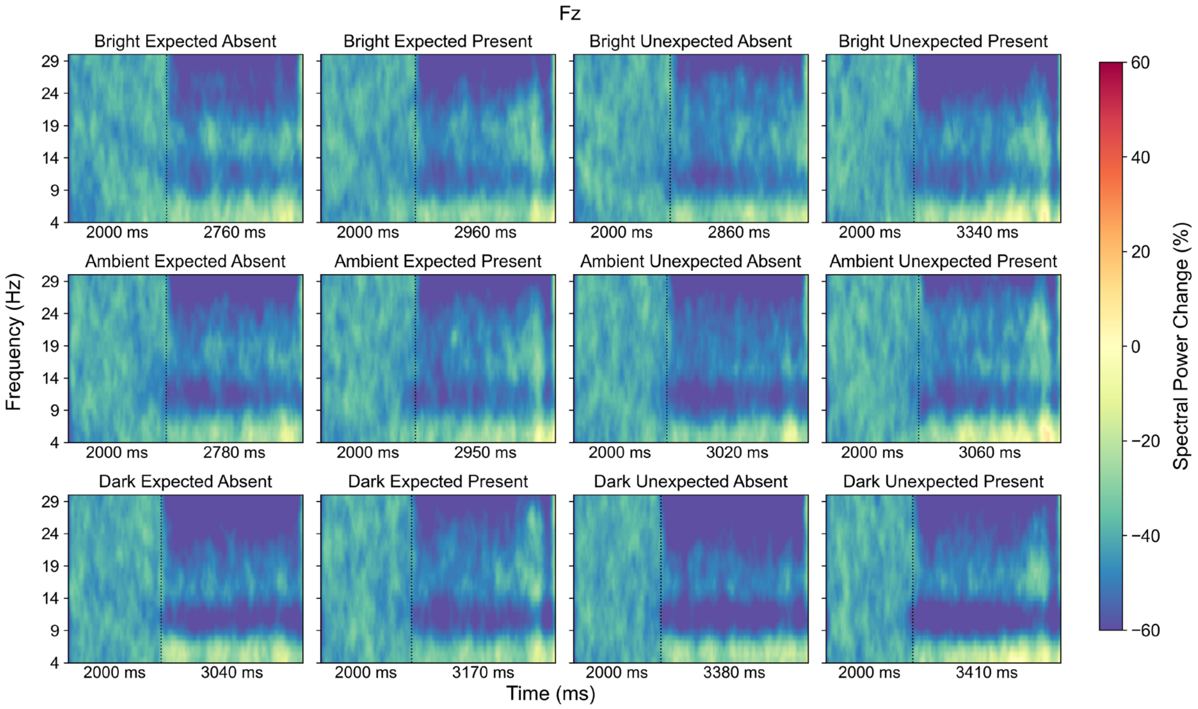


**A**

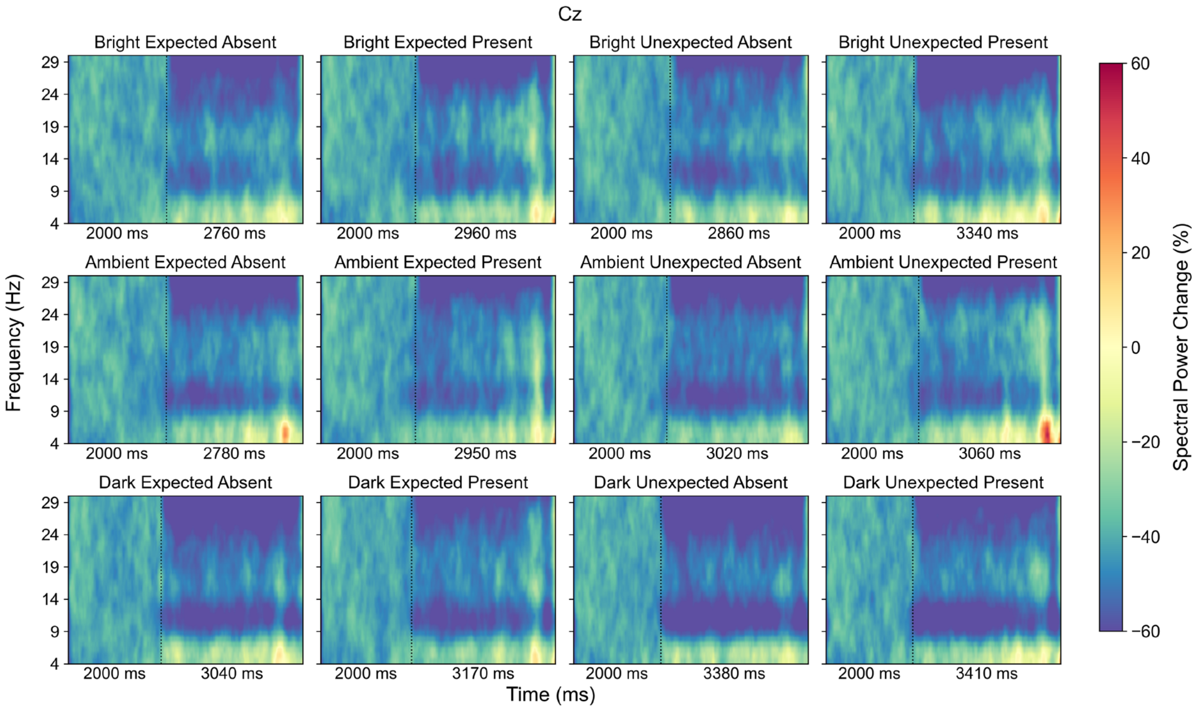


**B**

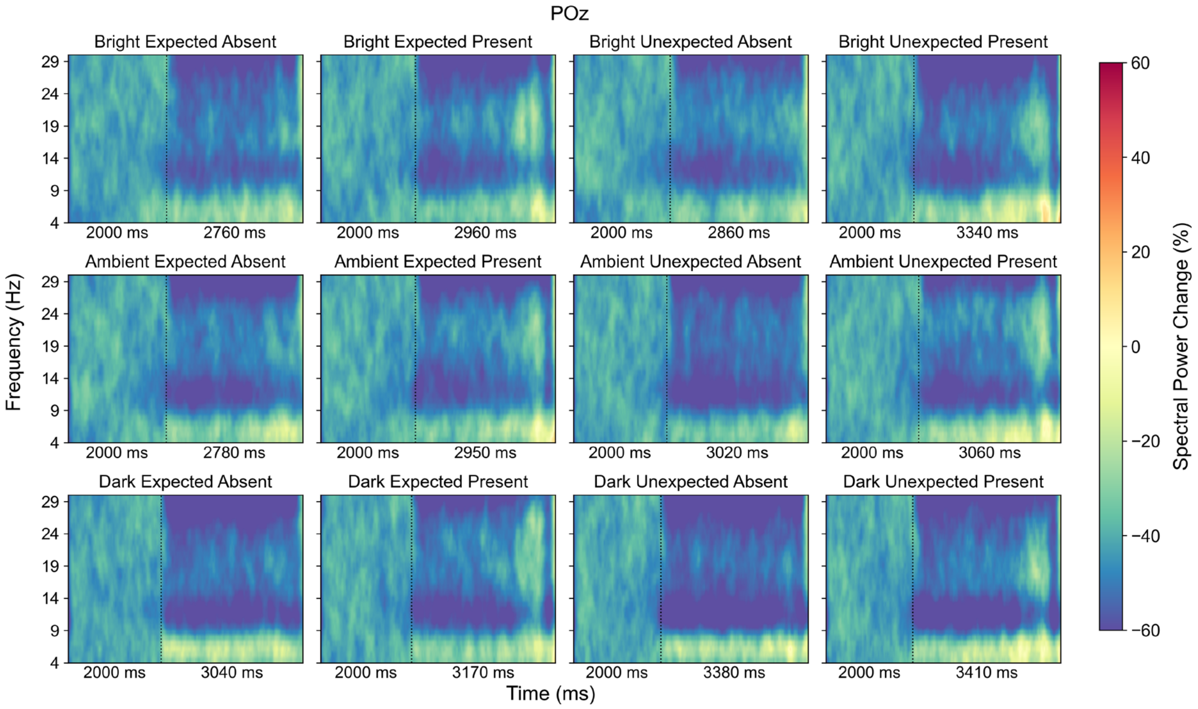


C

**Figure S1. TFRs at electrodes (A) Fz, (B) Cz, and (C) POz for the preparation phase. Frequencies range from 4–30 Hz. Each plot shows a 2s standing baseline period (left of the vertical dashed line) followed by the time-warped preparation phase (duration shown in ms). Values represent spectral power change (%) relative to baseline, with red colours indicating an increase in power and blue colours indicating a decrease in power.**

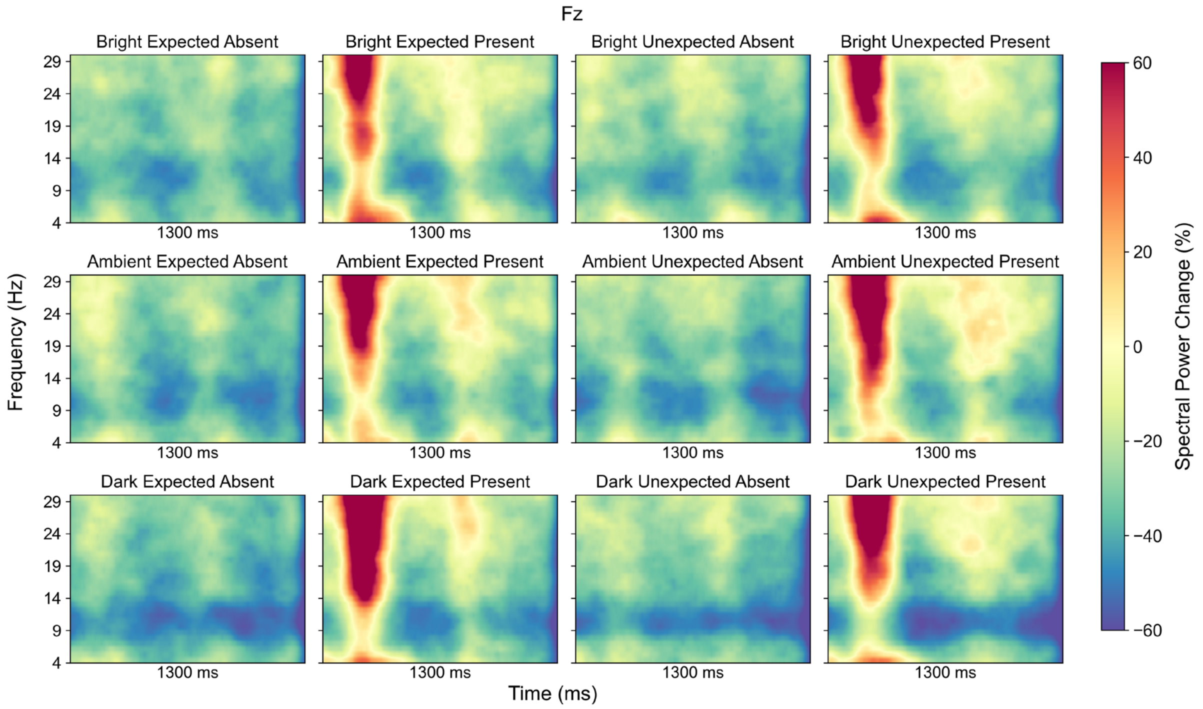

Fz
Bright Expected Absent
Bright Expected Present
Bright Unexpected Absent
Bright Unexpected Present
Ambient Expected Absent
Ambient Expected Present
Ambient Unexpected Absent
Ambient Unexpected Present
Dark Expected Absent
Dark Expected Present
Dark Unexpected Absent
Dark Unexpected Present
Frequency (Hz)
29
24
19
14
9
4
1300 ms
Time (ms)
Spectral Power Change (%)
60
40
20
0
−20
−40
−60


**A**

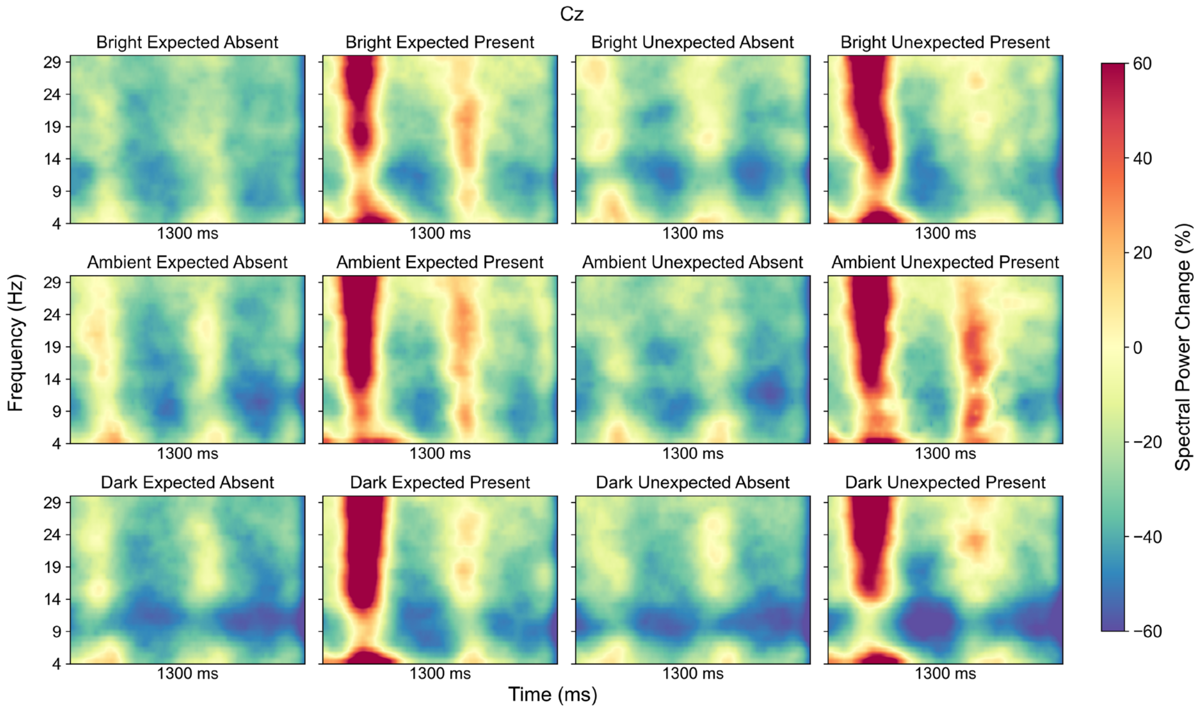

Cz
Bright Expected Absent
Bright Expected Present
Bright Unexpected Absent
Bright Unexpected Present
Ambient Expected Absent
Ambient Expected Present
Ambient Unexpected Absent
Ambient Unexpected Present
Dark Expected Absent
Dark Expected Present
Dark Unexpected Absent
Dark Unexpected Present
Frequency (Hz)
29
24
19
14
9
4
1300 ms
Time (ms)
Spectral Power Change (%)
60
40
20
0
−20
−40
−60


**B**

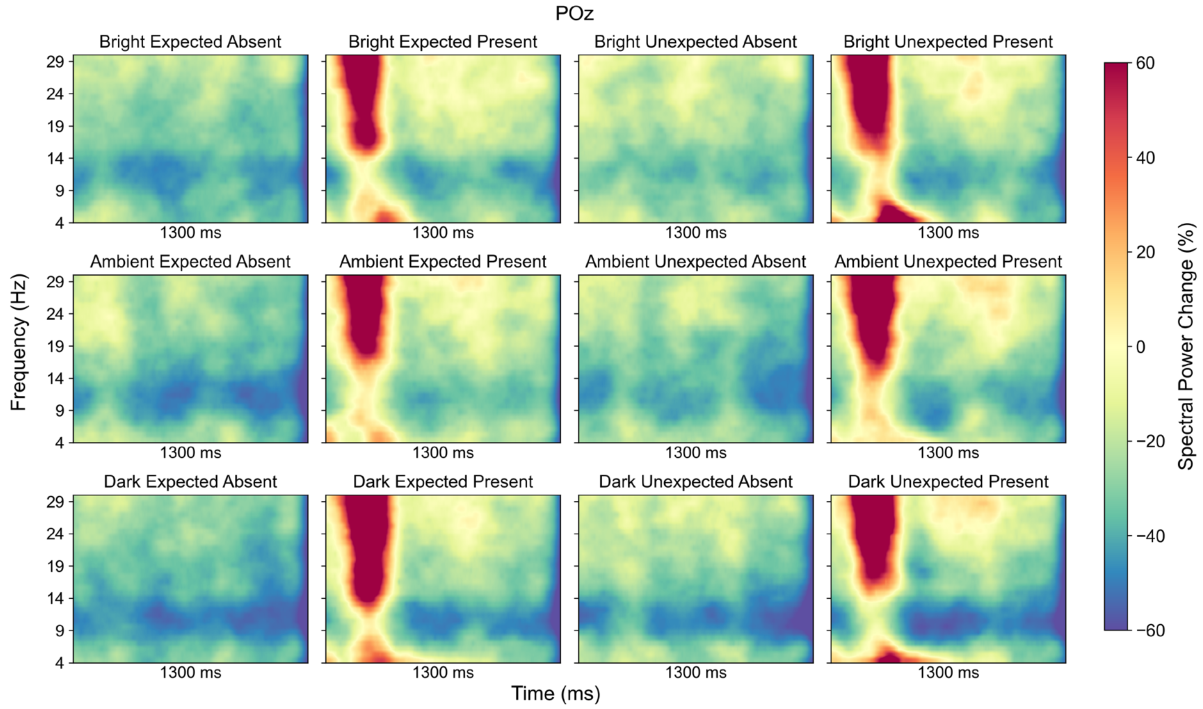


C

**Figure S2. TFRs at electrodes (A) Fz, (B) Cz, and (C) POz for the reset phase. Frequencies range from 4–30 Hz. Values represent spectral power change (%) relative to baseline, with red colours indicating an increase in power and blue colours indicating a decrease in power.**

#### S4. Region of interest analysis

A mixed ANOVA revealed no significant Age × Lighting interaction for frontal theta power $F(1,40) = 3.71$, $p = .061$, $\eta_p^2 = .09$. Similarly, no significant Age × Lighting interaction was found for occipito-parietal alpha power $F(1,40) = 0.74$, $p = .396$, $\eta_p^2 = .02$, or fronto-central beta power $F(1,40) = 0.03$, $p = .862$, $\eta_p^2 = .00$.

A paired t-test comparing Bright Expected Present and Dark Expected Present conditions for frontal theta power revealed no significant difference, $t(41) = -0.13$, $p = .90$, $d = 0.02$. Similarly, a paired t-test comparing Bright Unexpected Present and Dark Unexpected Present conditions for frontal theta power revealed no significant difference, $t(41) = 0.16$, $p = .876$, $d = 0.02$.

#### S5. Mean gait speed and cadence

**Table S2**

*Mean (standard error) gait speed and cadence for Age, Lighting, and Obstacle conditions.*

| Condition | Level | Speed (m/s) | Cadence (steps/min) |
|---|---|---|---|
| Age | Young Adults | 1.06 (0.02) | 100.94 (1.57) |
| | Older Adults | 0.93 (0.02) | 97.10 (1.69) |
| Lighting | Bright | 1.02 (0.02) | 99.61 (1.17) |
| | Ambient | 1.00 (0.02) | 99.46 (1.17) |
| | Dark | 0.97 (0.02) | 97.98 (1.17) |
| Obstacle presence | Present | 1.01 (0.02) | 97.75 (1.16) |
| | Absent | 0.99 (0.02) | 100.29 (1.16) |
| Obstacle × Expectation | Expected Absent | 1.01 (0.02) | 101.39 (1.17) |
| | Expected Present | 1.03 (0.02) | 98.56 (1.17) |
| | Unexpected Absent | 0.96 (0.02) | 99.19 (1.20) |
| | Unexpected Present | 0.99 (0.02) | 96.93 (1.20) |
